# Implicit anticipation of probabilistic regularities: Larger CNV emerges for unpredictable events

**DOI:** 10.1101/2020.08.14.251686

**Authors:** Andrea Kóbor, Zsófia Kardos, Kata Horváth, Karolina Janacsek, Ádám Takács, Valéria Csépe, Dezso Nemeth

## Abstract

Anticipation of upcoming events plays a crucial role in automatic behaviors. It is, however, still unclear whether the event-related brain potential (ERP) markers of anticipation could track the implicit acquisition of probabilistic regularities that can be considered as building blocks of automatic behaviors. Therefore, in a four-choice reaction time (RT) task performed by young adults (*N* = 36), the contingent negative variation (CNV) as an ERP marker of anticipation was measured from the onset of a cue stimulus until the presentation of a target stimulus. Due to the probability structure of the task, target stimuli were either predictable or unpredictable, but this was unknown to participants. The cue did not contain predictive information on the upcoming target. Results showed that the CNV amplitude during response preparation was larger before the unpredictable than before the predictable target stimuli. In addition, although RTs increased, the P3 amplitude decreased for the unpredictable as compared with the predictable target stimuli, possibly due to the stronger response preparation that preceded stimulus presentation. These results suggest that enhanced attentional resources are allocated to the implicit anticipation and processing of unpredictable events. This might originate from the formation of internal models on the probabilistic regularities of the stimulus stream, which primarily facilitates the processing of predictable events. Overall, we provide ERP evidence that supports the role of implicit anticipation and predictive processes in the acquisition of probabilistic regularities.

## 1. Introduction

Predicting future events is a core aspect of adaptive behavior. Predictions are guided by prior probabilities according to many theories of cognition, learning, and decision making (Daw, Gershman, Seymour, Dayan, & Dolan, 2011; Friston, 2005, 2010; Friston, Stephan, Montague, & Dolan, 2014; Gómez & Flores, 2011; Griffiths, Kemp, & Tenenbaum, 2008; Shohamy & Daw, 2015). Based on prior probabilities, the set of processes that facilitate perceptual, cognitive, and motor operations before their actual occurrence can be captured by the notion of anticipation (Hommel, 2009; van Boxtel & Böcker, 2004). Anticipation largely depends on the extraction and processing of probabilistic regularities organizing environmental events (e.g., Koelsch, Busch, Jentschke, & Rohrmeier, 2016; Maheu, Dehaene, & Meyniel, 2019; Meyniel, Maheu, & Dehaene, 2016), which have been found to be crucial in many automatized cognitive abilities (Armstrong, Frost, & Christiansen, 2017; Aslin, 2017; Conway, 2020). The presence and operation of anticipatory processes can be evidenced by reaction time (RT) changes and various measures derived from eye-tracking, electromyography, neuroimaging, and electrophysiology (e.g., Fan et al., 2007; Gómez & Flores, 2011; Killikelly & Szűcs, 2013; Medimorec, Milin, & Divjak, 2019; Tremblay & Saint-Aubin, 2009). Meanwhile, the question remains to what extent the electrophysiological markers of anticipation are sensitive to the processing of probabilistic regularities (Daltrozzo & Conway, 2014). Therefore, we investigated whether event-related brain potentials (ERPs) reflect the implicit anticipation of predictable and unpredictable regularities underlying the ongoing sensorimotor stream.

A possible candidate among the slow cortical potentials related to anticipatory processes is the contingent negative variation (CNV). The CNV has usually been recorded in cued/forewarned RT tasks as an increasingly negative potential shift occurring over the frontal and central electrode sites in the time interval between a cue/warning/S1 stimulus and a target/imperative/S2 stimulus requiring a response (Berchicci, Spinelli, & Di Russo, 2016; Brunia, 2003; Killikelly & Szűcs, 2013; Kononowicz & Penney, 2016; Leuthold, Sommer, & Ulrich, 2004; Molnár et al., 2008; Verleger, Paulick, Möcks, Smith, & Keller, 2013; Walter, Cooper, Aldridge, McCallum, & Winter, 1964). The CNV could be decomposed into multiple potentials: While the early phase of the component has been thought to reflect the orientation to the cue stimulus, the late phase could reflect the anticipatory activity for the target and the motor preparation of the related response (Loveless & Sanford, 1974; van Boxtel & Böcker, 2004; Weerts & Lang, 1973). The present study focuses on the late CNV, considering that the cue stimulus is not predictive of the target and a relatively short cue-target interval is used (see Stimuli, task, and procedure section) that might not ensure the full development of the different composing waves (Berchicci et al., 2016; Di Russo et al., 2017).

Given the different composing waves and the various paradigms applied, the suggested functional role of the CNV and its interpretation are also numerous. Beyond the above-listed processes and the original suggestion that the CNV indicates stimulus expectancy (Walter et al., 1964), this component has been linked to, for instance, time processing (Macar & Vidal, 2004), proactive control (Killikelly & Szűcs, 2013), motor pre/re-programming (Anatürk & Jentzsch, 2015; Jentzsch, Leuthold, & Richard ridderinkhof, 2004; Leuthold & Jentzsch, 2002), the complexity of preparation (Cui et al., 2000; De Kleine & Van der Lubbe, 2011), feedback/reward anticipation (Hackley, Valle-Inclán, Masaki, & Hebert, 2014), and the amount of attentional resources or effort recruited and available for stimulus processing and response preparation (Cui et al., 2000; Emerson, Daltrozzo, & Conway, 2014; Pauletti et al., 2014; Stadler, Klimesch, Pouthas, & Ragot, 2006). Overall, the CNV is likely a marker of a task-specific preparatory state that tunes and optimizes perceptual, cognitive, and motor processes, supported by the activation of multiple sensory-motor brain areas and controlled by the frontoparietal networks (Gómez & Flores, 2011; Kononowicz & Penney, 2016; Pauletti et al., 2014).

So far, only a few studies have examined directly the sensitivity of the CNV to the probabilistic regularities underlying a stimulus sequence while implicitly acquiring the sequence per se (Daltrozzo & Conway, 2014). In a visual S1-S2 associative learning task, the CNV amplitude decreased before the S2 in the predictable condition while it increased in the unpredictable condition as the acquisition of the probability structure progressed (Rose, Verleger, & Wascher, 2001). Meanwhile, the CNV amplitude increased for regular vs. random targets in an auditory learning-oddball task (Jongsma et al., 2006) and for possible vs. impossible target occurrence in another oddball task with tone pairs (Stadler et al., 2006). A further auditory sequential learning task showed that the CNV emerged only after the high- and low-probability predictor tones and not after the zero-probability predictors (Emerson et al., 2014). Overall, it seems that either enhanced or decreased CNV amplitudes could indicate the sensitivity to probabilistic regularities.

In this study, therefore, we aimed to measure implicit anticipatory processes in a four-choice RT task that, unknown to participants, included a sequential regularity between non-adjacent trials yielding a probability structure with predictable and unpredictable stimuli. By means of RTs, using a version of this task, we have already shown that the implicitly acquired prior knowledge on the probabilistic regularities influenced the processing of further stimuli lacking a predictable structure (Kóbor, Horváth, Kardos, Nemeth, & Janacsek, 2020). We assumed that this persistence of knowledge occurred through the formation of internal models. Therefore, it is conceivable that anticipatory processes based on similar internal models would start to operate before each stimulus is presented, at least after the probability structure of the task has been acquired. Because of these internal models, the anticipation of predictable stimuli and the preparation of appropriate responses possibly become automatic and require less attentional resources, as opposed to the unpredictable stimuli (Horváth et al., in press). Accordingly, considering the functional significance of the CNV, its amplitude was expected to be larger for unpredictable than for predictable stimuli. In parallel, according to other behavioral studies with this task (e.g., D. V. Howard et al., 2004; Nemeth et al., 2010; Takács et al., 2018; Tóth et al., 2017), we expected slower RTs to unpredictable than to predictable stimuli.

## 2. Material and methods

### 2.1 Participants

Thirty-six healthy young adults (23 females) between the ages of 18 and 28 (*M* = 21.6, *SD* = 2.2) participated in the study. They were undergraduate students from Budapest, Hungary (years of education: *M* = 14.6, *SD* = 1.5). Handedness was assessed with the Edinburgh Handedness Inventory revised version (Dragovic, 2004a, 2004b; Oldfield, 1971), according to which the mean Laterality Quotient was 80.3 (*SD* = 35.3; −100 means complete left-handedness, 100 means complete right-handedness). Participants had normal or corrected-to-normal vision, and according to the pre-defined inclusion criteria, none of them reported a history of any neurological and/or psychiatric condition, and none of them was taking any psychoactive medication. They performed in the normal range on standard neuropsychological tests that were administered before the EEG experiment, part of the standard participant screening procedure we follow in the lab (Wisconsin Card Sorting Task [perseverative error percentage]: *M* = 10.42, *SD* = 2.23; Digit span task [mean short-term memory span; possible range: 3–9]: *M* = 6.31, *SD* = 1.31; Counting span task [mean working memory span; possible range: 2–6]: *M* = 3.64, *SD* = 0.72; Go/No-Go task [discriminability index: hit rate minus false alarm rate]: *M* = .76, *SD* = .15; these results are not published elsewhere). All participants provided written informed consent before enrollment and received course credit for taking part in the study. The study was approved by the United Ethical Review Committee for Research in Psychology (EPKEB) in Hungary and was conducted in accordance with the Declaration of Helsinki.

### 2.2 Stimuli, task, and procedure

Implicit acquisition of probabilistic regularities was measured by a modified cued version of the Alternating Serial Reaction Time (ASRT) task (Horváth et al., in press; Kóbor et al., 2019; Kóbor et al., 2018; Nemeth et al., 2010). In this task version, an experimental trial consisted of the presentation of two subsequent stimuli: a cue and a target. First, a central fixation cross that served as the cue was presented for a fixed duration of 900 ms. Second, after this duration, the cue was replaced by an arrow stimulus that served as the target, presented at the center of the screen for a fixed duration of 200 ms (cf. Kóbor et al., 2019; Kóbor et al., 2018). After target offset, a blank response window was displayed for a fixed duration of 700 ms. Then, the next trial started, yielding an 1800-ms-long intertrial interval (ITI) that consisted of an anticipatory/response preparation phase (from cue onset to target onset) and a response execution phase (from target onset to the next cue onset, see Fig. 1A).

**Figure 1.**
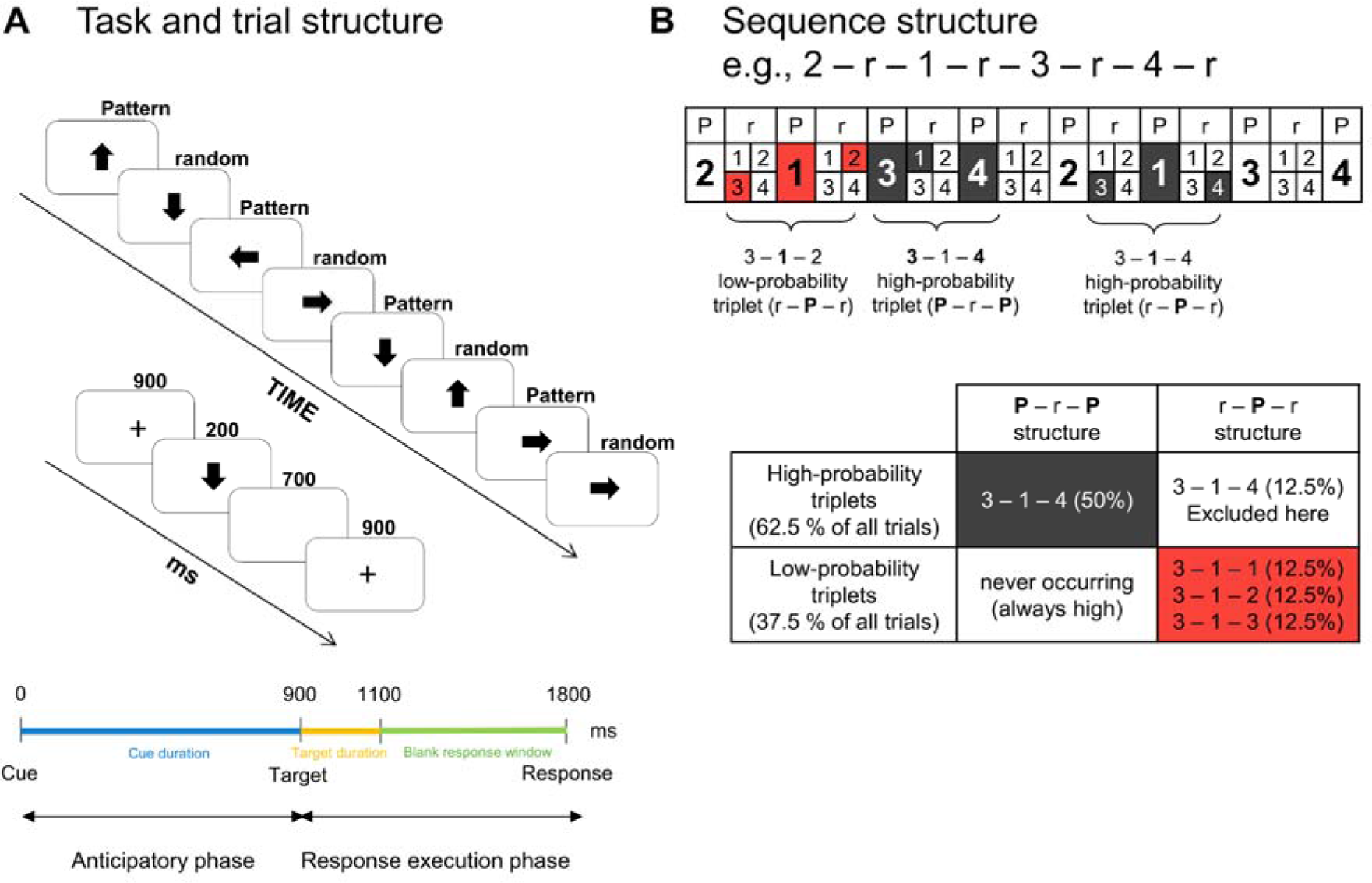
Design of the experiment. (A) In this version of the Alternating Serial Reaction Time (ASRT) task, a central fixation cross that serves as the cue is followed by an arrow stimulus that serves as the target. The cue is not predictive of the target. The duration of the cue is 900 ms, the duration of the target is 200 ms, and responses are recorded from target onset until the end of the trial (for altogether 900 ms). This yields a fixed 1800-ms-long intertrial interval (ITI). The presentation of the target stimuli follows an eight-element sequence, within which pattern (P) and random (r) elements alternate with one another. (B) In the alternating sequence structure, numbers denote the four spatial directions (1 = left, 2 = up, 3 = down, 4 = right) of the arrows. The alternating sequence makes some runs of three consecutive trials (triplets) more probable than others. High-probability triplets are denoted with dark-grey shading and low-probability triplets are denoted with coral shading. Each pattern and random trials are categorized as the last trial of a high- or a low-probability triplet, thus, three different triplets occur: pattern high-probability (P – r – P structure), random high-probability (r – P – r structure), and random low-probability (r – P – r structure) triplets. This study analyses only pattern high-probability or predictable (dark-grey shading in the lower table) and random low-probability or unpredictable triplets (coral shading in the lower table).

Participants were instructed to respond as quickly and accurately as possible to the direction of the target by pressing one of the four response keys of a Cedrus RB-530 response pad (Cedrus Corporation, San Pedro, CA). The spatial directions of the arrow stimuli were mapped onto the four response keys (left direction/left response key = left thumb, up = left index finger, down = right thumb, and right = right index finger). Participants were also asked to pay attention to and maintain their gaze on the fixation cross and try not to blink, because it was shortly followed by the target. If an incorrect response occurred, the fixation cross starting the next trial was presented in red color instead of the original black one. If no response occurred during target presentation or in the predefined response window (i.e., missing response), the fixation cross starting the next trial was presented in blue instead of black. Participants were informed that the color of the fixation cross would change if they did not provide the correct response. As the fixation cross did not convey information on the *following* target, no other details were provided about its function. After an incorrect or missing response, although participants could provide further behavioral responses, this did not influence the presentation and timing of the next trial.

Unbeknownst to the participants, the arrow stimuli were presented according to an eight-element sequence, within which predetermined/pattern (P) and random (r) elements alternated with one another (see Fig. 1A). For instance, 2 – r – 1 – r – 3 – r – 4 – r was one of the sequences, where numbers denoted the four predetermined spatial directions [1 = left, 2 = up, 3 = down, 4 = right] of the arrows and *r*s denoted the randomly chosen directions out of the four possible ones (see Fig. 1B). There were 24 permutations of the four spatial directions that could determine the applied sequence; however, because of the continuous presentation of the stimuli, there were only six *unique* permutations: 1 – r – 2 – r – 3 – r – 4 – r, 1 – r – 2 – r – 4 – r – 3 – r, 1 – r – 3 – r – 2 – r – 4 – r, 1 – r – 3 – r – 4 – r – 2 – r, 1 – r – 4 – r – 2 – r – 3 – r, and 1 – r – 4 – r – 3 – r – 2 – r. Note that each of these six permutations could start at any position; i.e., 1 – r – 3 – r – 4 – r – 2 – r and 2 – r – 1 – r – 3 – r – 4 – r were identical sequence permutations. The applied sequence for each participant was determined by *only one* of the six unique permutations of the 24 possible ones. The given permutation was selected for each participant in a pseudorandom manner (see also J. H. Howard, Jr. & Howard, 1997; Kóbor et al., 2019; Kóbor et al., 2018; Nemeth et al., 2010).

One block of the ASRT task contained 85 trials. In each block, the eight-element sequence repeated 10 times after five warm-up trials consisting only of random stimuli. The trials of each block were categorized as chunks of three successive trials, hereafter referred to as *triplets*. Particularly, each trial was categorized as the third trial of a triplet and the third trial of a triplet was also the second trial of the following triplet, e.g., 2 – 3 – ****1****, 3 – ****1**** – 2 (Kóbor, Janacsek, Takács, & Nemeth, 2017; Kóbor et al., 2018; Szegedi-Hallgató et al., 2017). The warm-up trials at the beginning of each block were not categorized as triplets. The warm-up trials were followed by two starter trials containing the first two elements of the alternating sequence. These two starter trials were not categorized as triplets either, since also the foremost triplet technically requires three successive trials. Thus, trials were categorized as triplets from the 8^th^ trial of the block. Altogether 30 blocks, containing 2550 trials in total, were completed. From these trials, 2340 triplets were constructed, but not all of them were used in the analysis (see Behavioral data analysis section below).

The alternating sequence yields a probability structure in which some triplets are more probable than others. Thus, the construction of triplets could be considered as a method for identifying a hidden probability structure in the ASRT task. Particularly, while the third trials of some triplets are *probable* (predictable) continuations for the first trials, the third trials of other triplets are *less probable* continuations for the first trials. The former triplets are referred to as *high-probability* triplets while the latter ones are referred to as *low-probability* triplets (e.g., Nemeth & Janacsek, 2011; Nemeth, Janacsek, Polner, & Kovacs, 2013). Because of the alternating sequence, although high-probability triplets could have P – r – P or r – P – r structure, low-probability triplets could only have a r – P – r structure (see Fig. 1B). In the case of the 2 – r – 1 – r – 3 – r – 4 – r sequence, 2 – X – 1, 1 – X – 3, 3 – X – 4, and 4 – X – 2 are high-probability triplets (X denotes the middle trial of the triplet), and, for instance, 1 – X – 2 and 4 – X – 3 are low-probability ones. From another perspective, random trials that are the 50% of all trials appear either with high or low probability, while pattern trials that are the other 50% of all trials always appear with high probability. Overall, the combination of the sequential and probability properties yields three possible triplet types: pattern high-probability, random high-probability, and random low-probability triplets (occurring with an overall probability of 50%, 12.5%, and 37.5%, respectively; see Fig. 1B).

After completing each block, participants received feedback (lasting for 4000 ms) about their mean reaction time and accuracy in the given block, then, they could have a short rest before starting the next block. The experimental procedure lasted about 2.5 hours, including the application and removal of the electrode cap. The ASRT task was written in and controlled by the Presentation software (v. 18.1, Neurobehavioral Systems). Stimuli were displayed on a 21” LCD screen at a viewing distance of 125 cm. Neuropsychological tests (see Participants section) were administered a few days before the EEG experiment during a one-hour-long session.

### 2.3 Behavioral data analysis

The set of the constructed triplets was narrowed for further analysis. First, following the standard data analysis protocol established in previous studies using the ASRT task (e.g., J. H. Howard, Jr. & Howard, 1997; Kóbor et al., 2017; Nemeth, Janacsek, Polner, et al., 2013; Song, Howard, & Howard, 2007; Virag et al., 2015), two types of low-probability triplets – repetitions (e.g., 1 – 1 – 1, 4 – 4 – 4) and trills (e.g., 1 – 2 – 1, 2 – 4 – 2) – were excluded, because pre-existing response tendencies have often been shown to them (D. V. Howard et al., 2004). Therefore, the analyzed low-probability triplet category consisted of low-probability triplets without trills and repetitions (but see Szegedi-Hallgató, Janacsek, & Nemeth, 2019).

Second, all random high-probability triplets were excluded from further analyses (cf. Horváth et al., in press). Random high-probability triplets could be considered as “accidentally-regular” random triplets, which are predictable but rarely occurring at the level of unique triplets. These trials integrate characteristics related to both the alternating P – r – P – r sequence and the hidden probability structure (Kóbor et al., 2019). In addition, some pre-existing response biases originating from higher levels of the alternating sequence have also been observed for these triplets (Szegedi-Hallgató et al., 2019). Therefore, the exclusion of random high-probability triplets enabled us to focus on contrasting the predictable pattern events with the unpredictable random events (i.e., pattern high-probability triplets vs. random low-probability triplets),

Ten-block-long bins of the behavioral data were grouped into three periods, labeled consecutively in this paper (1, 2, etc.). For each participant and period, separately for pattern high-probability and random low-probability triplets, median RT was calculated for those correct responses that also followed correct responses. Independent of triplet types, general skill improvements (faster RTs) reflecting more efficient visuomotor and motor-motor coordination due to practice were also considered (cf. Hallgató, Győri-Dani, Pekár, Janacsek, & Nemeth, 2013; Juhasz, Nemeth, & Janacsek, 2019).

The mean accuracy of responding for each triplet type and period are provided in Table 1; otherwise, the paper focuses on the RT analysis (cf. Kóbor et al., 2019; Kóbor et al., 2018). We have three reasons to do so: accuracy is influenced by (1) the feedback given to participants after each block; (2) the presentation of the cue increased the length of the trial that possibly provided more time for accurate response selection; and (3) overall accuracy has usually been high with relatively low variability in samples of healthy young adults performing the ASRT task (J. H. Howard, Jr. & Howard, 1997; Janacsek, Ambrus, Paulus, Antal, & Nemeth, 2015; Nemeth et al., 2010; Romano, Howard, & Howard, 2010).

**Table 1.** Mean percentage (%) and standard deviation of response accuracy split by triplet type and period.

|  | Pattern High | Random Low |
| --- | --- | --- |
|  | *M (SD)* | *M (SD)* |
| Period<sub>1</sub> | 96.0 (2.8) | 95.8 (3.0) |
| Period<sub>2</sub> | 95.9 (2.8) | 95.6 (2.5) |
| Period<sub>3</sub> | 95.8 (2.6) | 95.1 (3.2) |
| Overall | 95.9 (2.5) | 95.5 (2.6) |

### 2.4 EEG recording and analysis

The continuous EEG activity was recorded in an electrically shielded, acoustically attenuated, and dimly lit room using the actiCAP active electrode system with BrainAmp Standard amplifier and BrainVision Recorder 1.2 software (BrainProducts GmbH, Gilching, Germany). The 64 sensors consisting of Ag/AgCl electrodes integrated with active circuits were mounted in an elastic cap and placed according to the 10% equidistant system. The FCz electrode was used as reference and the Fpz electrode was used as ground. The sampling rate was 1000 Hz. During recording, the impedance of the electrodes was kept below 10 kΩ.

The continuous EEG data were analyzed offline using the BrainVision Analyzer 2.1.2 software (BrainProducts GmbH). The pre-processing steps described below followed those presented in the Kóbor et al. (2018) and Kóbor et al. (2019) papers with minor modifications (e.g., filter settings, length of the segment, number of time bins) to fit the purposes of the present study. First, after visual screening for major deflections, if necessary, bad electrodes were replaced by spline interpolation: A maximum of one electrode per participant (*M* = 0.14, *SD* = 0.35) was interpolated. Second, the EEG data were band-pass filtered within 0.03-30 Hz (48 dB/oct) and notch filtered at 50 Hz to remove additional electrical noise. Third, horizontal and vertical eye-movement artifacts and heartbeats were corrected with independent component analysis (Delorme, Sejnowski, & Makeig, 2007): Components between one and three per participant (*M* = 1.97, *SD* = 0.51) were rejected, then, the channel-based EEG data were recomposed. Fourth, EEG data were re-referenced to the average activity of all electrodes. (The FCz, which was the online reference electrode, was also included in the calculation of the offline average reference and then re-used to analyze its signals, as well.) Fifth, the continuous EEG data were segmented in two steps as follows.

To track the temporal trajectory of acquisition, as in the case of the behavioral data, the data were cut into three periods, each containing ten consecutive blocks of the ASRT task. Next, within each period, segments were extracted from −200 to 1500 ms relative to cue onset, separately for pattern high-probability and random low-probability triplets. Only those correctly responded triplets were included in this step of the segmentation that also followed correctly responded triplets. Triplets with any responses occurring during the response preparation phase (i.e., before the presentation of the target) were omitted from the analyses. As in the case of the behavioral data, trills and repetitions were also excluded and random high-probability triplets were not analyzed. Altogether six (two triplet types * three periods) segment types were created for the cue-locked averages.

Following segmentation, the segments were baseline corrected based on the mean activity from −200 ms to 0 ms (pre-cue baseline). Then, to remove artifacts still present in the data after ICA corrections, an automatic artifact rejection algorithm implemented in the BrainVision Analyzer software was applied, which rejected segments where the activity exceeded +/−100 μV at any of the electrode sites. In addition, after this automated cleaning procedure, the retained segments were visually inspected for low frequency electrode drifts that could confound the CNV measurement. Segments with low frequency electrode drifts were manually rejected. After, the mean numbers of the retained (artifact-free) segments were 336.0 (*SD* = 43.2, range: 185 – 382) for pattern high-probability triplets and 167.1 (*SD* = 24.5, range: 75 – 210) for random low-probability triplets. This means that the mean proportion of the removed segments was 6.1% for pattern high-probability triplets and 6.3% for random low-probability triplets. Finally, the retained (artifact-free) segments were averaged for the two triplet types in each of the three periods.

Grand average ERP waveforms calculated separately for each triplet type in each period as well as averaged for the entire acquisition phase across all periods were created. These grand averages were visually inspected to determine the latency range where the CNV component would emerge and vary as a function of triplet types. Accordingly, the *time window* for the CNV analysis was determined in a post-hoc manner between 800 ms and 900 ms after cue onset (cf. Boehm, van Maanen, Forstmann, & van Rijn, 2014). For both triplet types, the CNV showed maximal negativity in this time window, after its steep increase starting around 700 ms following cue onset over the frontocentral, central, and centroparietal electrode sites. The *electrodes* for CNV analysis were also chosen in a post-hoc manner based on the difference waveforms calculated as ERPs for random low-probability triplets minus ERPs for pattern high-probability triplets. The difference waveforms were maximal over the central and centroparietal electrode sites (cf. Brunia & Damen, 1988; De Kleine & Van der Lubbe, 2011; Gómez, Marco, & Grau, 2003; Leynes, Allen, & Marsh, 1998; Verleger, Wauschkuhn, van der Lubbe, Jaśkowski, & Trillenberg, 2000). Therefore, based on the observed topographical distribution of the CNV difference waveform, a centroparietal (CP) electrode pool was defined by calculating the average activity of the electrodes C1, Cz, C2, CP1, CPz, CP2, P1, Pz, and P2. Altogether, the CNV was quantified over this centroparietal electrode pool as the mean amplitude between 800 ms and 900 ms after cue onset for all triplet types and periods.

The distribution of the between-triplets difference remained stable after target onset, which is in line with the observation that the CNV and the P3 components could overlap (Verleger, Paehge, Kolev, Yordanova, & Jaśkowski, 2006; Verleger et al., 2013; Verleger, Siller, Ouyang, & Śmigasiewicz, 2017). Therefore, the peak of the P3 and the late descending flank of the P3 (hereafter referred to as the late P3) were also examined over the same centroparietal electrode pool. The peak of the P3 was quantified as the mean amplitude between 1200 ms and 1300 ms after cue onset. The late P3 was quantified in the remaining interval of the segment, as the mean amplitude between 1300 ms and 1500 ms (cf. Kóbor et al., 2019).

### 2.5 Statistical analysis

The implicit acquisition and prediction of probabilistic regularities was quantified with two-way repeated measures analyses of variance (ANOVA) with Type (pattern high-probability vs. random low-probability triplet) and Period (1–3) as within-subjects factors on the RTs, the mean amplitude of the CNV, and the mean amplitudes of the P3 peak and the late P3. In all ANOVAs, the Greenhouse-Geisser epsilon (ε) correction (Greenhouse & Geisser, 1959) was used when necessary, i.e., when sphericity was violated as indicated by the significance of the Mauchly’s sphericity test. (The test statistics and the *p* values for the Mauchly’s test are not reported.) Note that the assumption of sphericity was not relevant in the case of the Type factor as it involved less than three levels. Original *df* values and corrected *p* values (if applicable) are reported together with partial eta-squared (η_*p*_^2^) as the measure of effect size. LSD (Least Significant Difference) tests for pairwise comparisons were used to control for Type I error.

## 3. Results

### 3.1 Behavioral results

The Type (pattern high-probability vs. random low-probability triplet) by Period (1–3) ANOVA on the RTs revealed a significant main effect of Type, *F*(1, 35) = 8.86, *p* = .005, η_*p*_^2^ = .202, indicating faster RTs on pattern high-probability triplets than on random low-probability ones (362.3 ms vs. 364.3 ms; see Fig. 2A). Although the Type * Period interaction was only a tendency, *F*(2, 70) = 2.42, *p* = .096, η_*p*_^2^ = .065, pairwise comparisons showed that the difference between pattern high-probability and random low-probability triplets was significant in period_3_ (3.5 ms, *p* = .004) but not before (period_1_: 1.4 ms, *p* = .144, period_2_: 1.0 ms, *p* = .217). In addition, this difference was significantly larger in period_3_ than in period_2_ (*p* = .026; other pairwise differences were nonsignificant, *p* ≥ .136). In details, RTs on pattern high-probability triplets tended to become faster as the task progressed (period_3_ vs. period_1_:360.4 ms vs. 364.7 ms, *p* = .052); meanwhile, the fastest RTs on low-probability triplets were found in period_2_ (period_2_ vs. period_1_: 362.7 ms vs. 366.1 ms, *p* = .028; other pairwise differences did not approach significance for both triplet types, *p* ≥ .120). General skills did not reliably improve: The Period main effect was only a tendency, *F*(2, 70) = 2.39, *p* = .099, η_*p*_^2^ = .064, and, according to the pairwise comparisons, improvement between period_1_ and period_2_ just reached significance (365.4 ms vs. 362.2, *p* = .050) and no further change was observed (*p*s ≥ .113). Overall, RTs differed between the triplet types and some indication was found for the stabilization of this difference by the end of the task.

**Figure 2.**
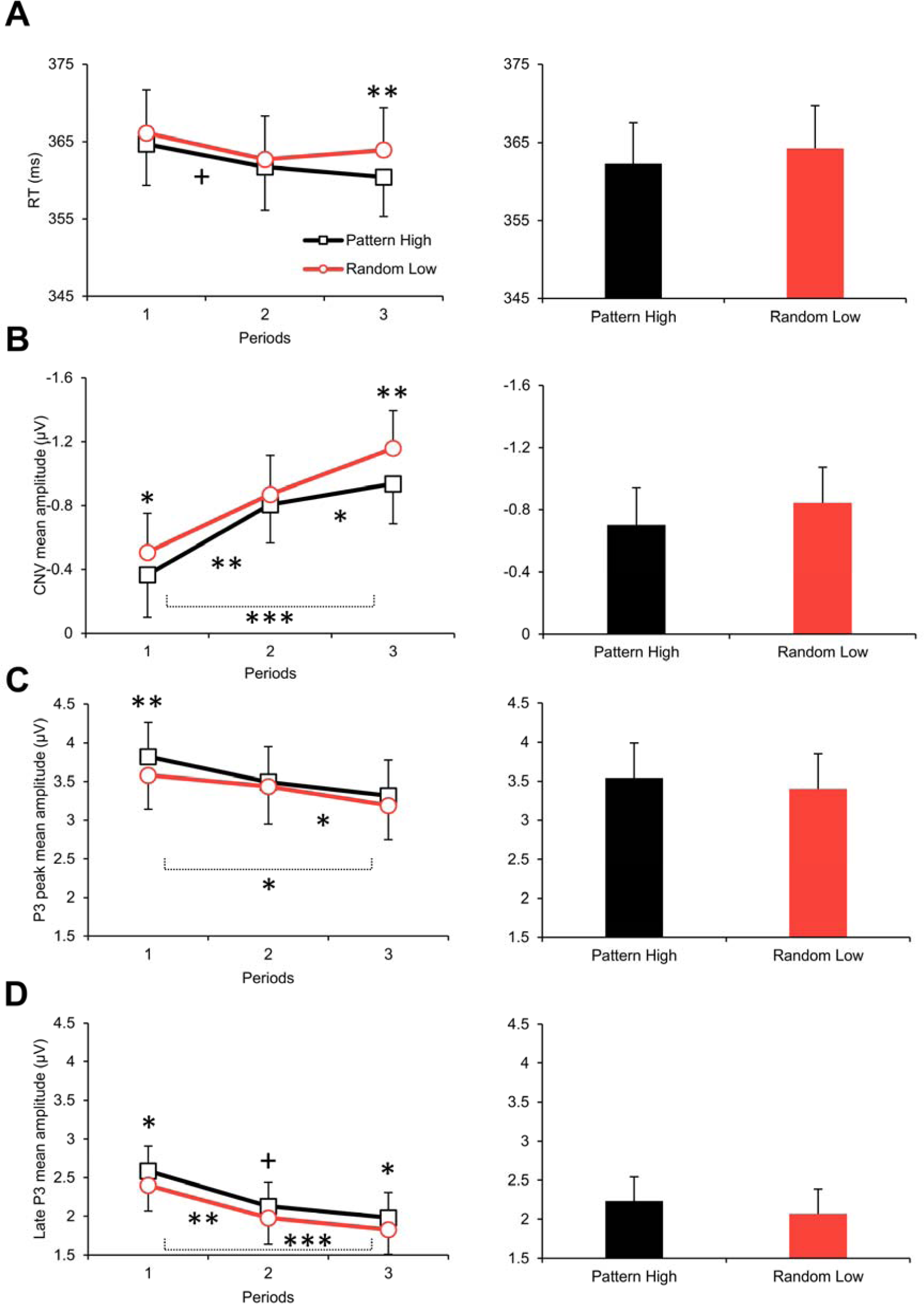
(A) Group-average RTs of correctly responded triplets split by period (1–3) and triplet type (pattern high-probability vs. random low-probability triplets) (left panel, line chart) and split by triplet type but collapsed across periods (right panel, bar chart). (B) Group-average CNV mean amplitudes split by period and triplet type (left panel, line chart) and split by triplet type but collapsed across periods (right panel, bar chart). Values along the vertical axes are plotted in a reversed scale. (C) Group-average P3 peak mean amplitudes split by period and triplet type (left panel, line chart) and split by triplet type but collapsed across periods (right panel, bar chart). (D) Group-average late P3 mean amplitudes split by period and triplet type (left panel, line chart) and split by triplet type but collapsed across periods (right panel, bar chart). Error bars denote standard error of mean. On the left panel, asterisks/plus marks above the means of each period denote the significance of the difference between triplet types (note that these pairwise differences are indicated here although nonsignificant Type * Period interactions were found). Asterisks/plus marks below the line charts denote the pairwise differences between periods collapsed across triplet types (i.e., pairwise differences related to the Period main effects). *Note*: + *p* < .010, * *p* < .050, ** *p* < .010, *** *p* < .001

### 3.2 ERP results

Grand average ERP waveforms for the two triplet types averaged for all periods and split by period over the centroparietal electrode pool are presented in Figures 3–4, respectively.

**Figure 3.**
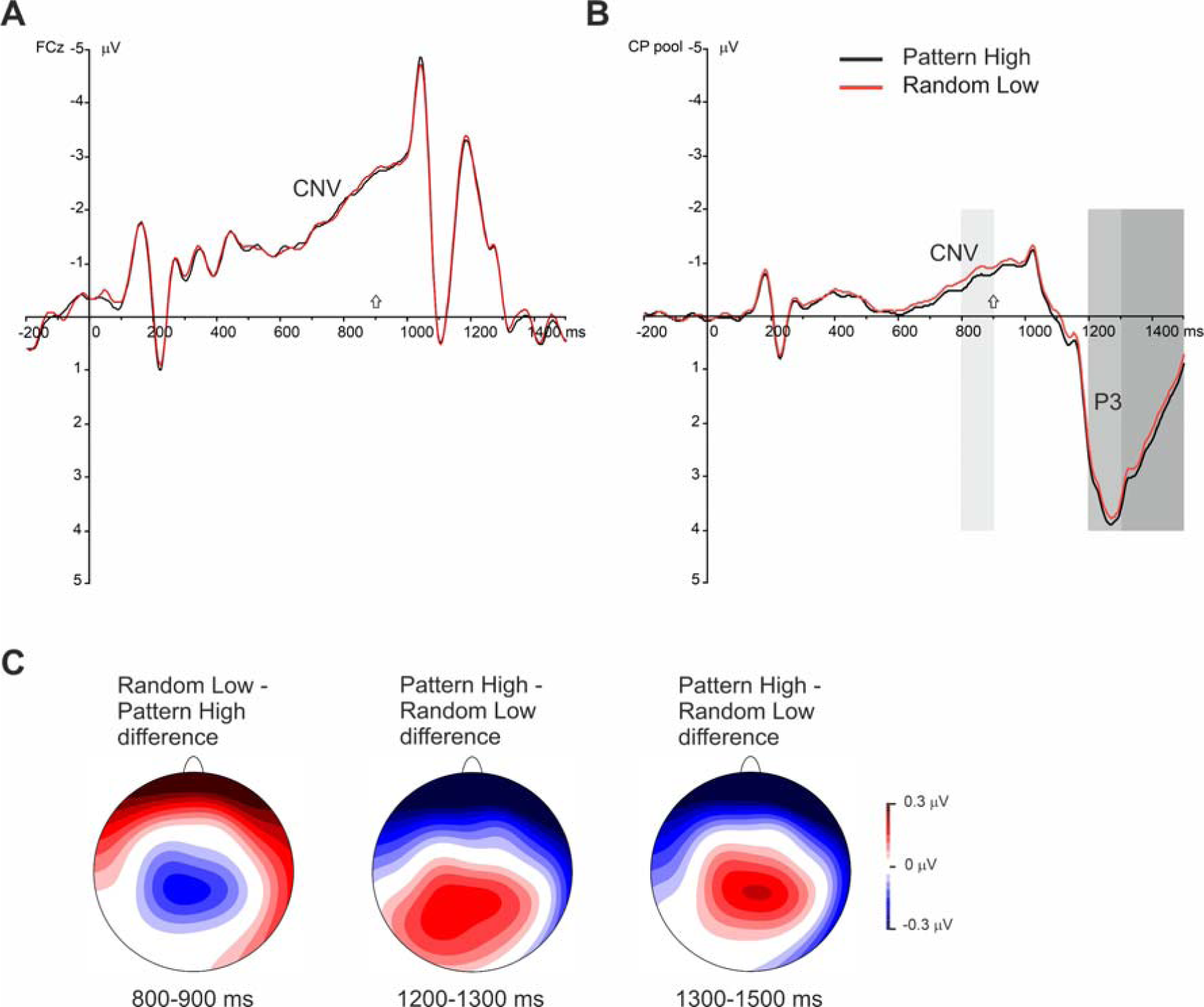
Grand average ERP waveforms at (A) electrode FCz and over (B) the centroparietal electrode pool are presented, displaying the CNV and the P3 components for pattern high-probability and random low-probability triplets, averaged for all periods. The cue onset was at 0 ms, the target onset was at 900 ms; the latter is denoted by small arrows above the horizontal axes. In part (B) of this figure, the light-grey shaded area indicates the time window in which the CNV was quantified (800-900 ms), the medium-grey shaded area indicates the time window in which the P3 peak was quantified (1200-1300 ms), and the dark-grey shaded area indicates the time window in which the late P3 was quantified (1300-1500 ms). Negativity is plotted upwards here and in Figure 4. (C) The scalp topography (amplitude distribution) of ERP differences for random low-probability minus pattern high-probability triplets and for pattern high-probability minus random low-probability triplets in the time windows of the CNV, the P3 peak, and the late P3, respectively (from left to right), averaged for all periods.

**Figure 4.**
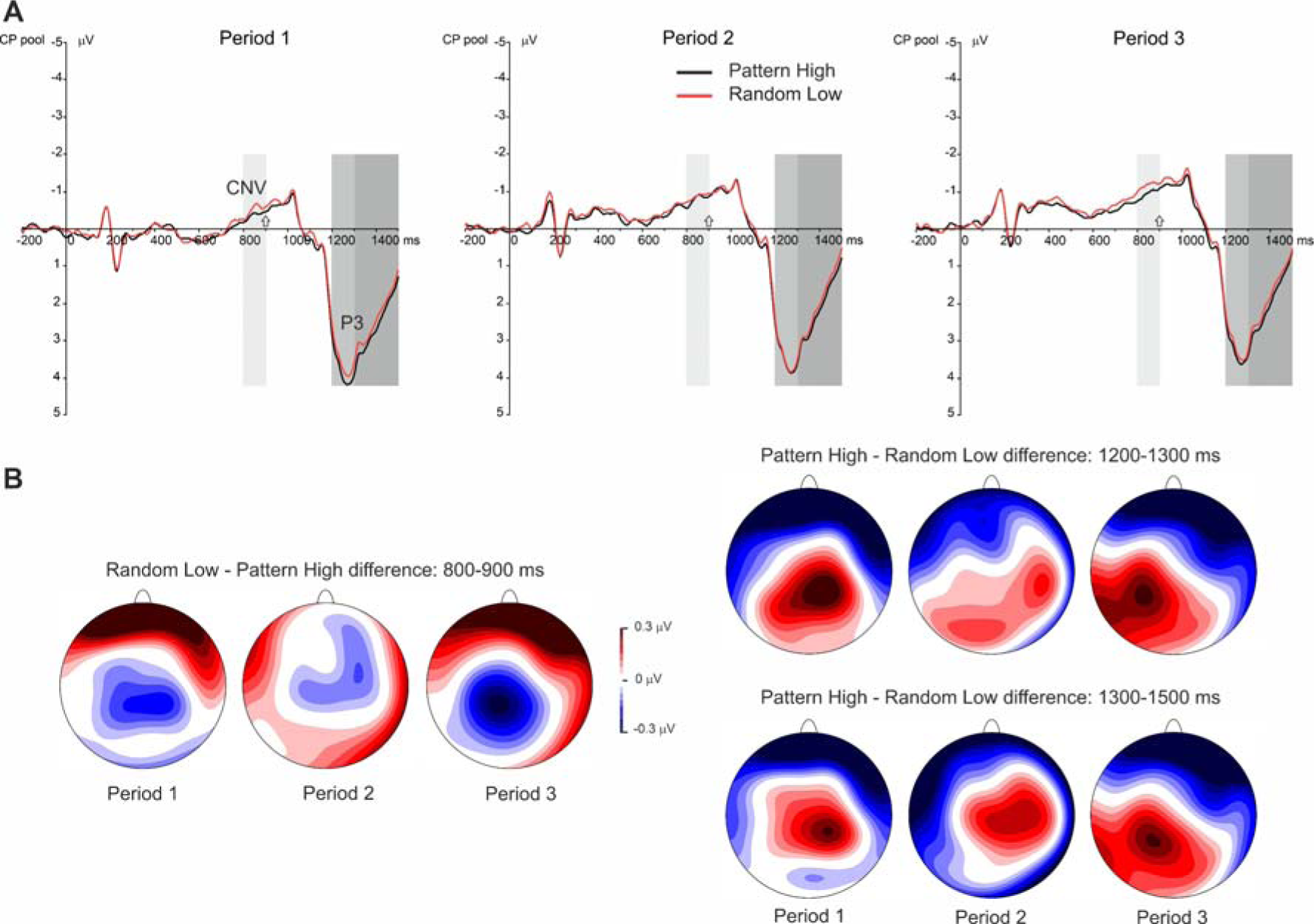
(A) Grand average ERP waveforms over the centroparietal electrode pool are presented, displaying the CNV and the P3 components for each period (1–3) and triplet type (pattern high-probability and random low-probability triplets). The cue onset was at 0 ms, the target onset was at 900 ms; the latter is denoted by small arrows above the horizontal axes. The light-grey shaded area indicates the time window in which the CNV was quantified (800-900 ms), the medium-grey shaded area is the time window of the P3 peak (1200-1300 ms), and the dark-grey shaded area is the time window of the late P3 (1300-1500 ms). (B) The scalp topography (amplitude distribution) of ERP differences in each period for random low-probability minus pattern high probability triplets in the time window of the CNV (left panel), and for pattern high-probability minus random low-probability triplets in the time windows of the P3 peak (right panel, top tow) and the late P3 (right panel, bottom row).

#### 3.2.1 CNV

The Type by Period ANOVA on the CNV showed a significant main effect of Type, *F*(1, 35) = 12.33, *p* = .001, η_*p*_^2^ = .261, indicating that the mean amplitude of the CNV was larger (more negative) for random low-probability triplets than for pattern high-probability ones (−0.84 μV vs. −0.70 μV; see Fig. 2B and Fig. 3B, C). This difference did not change with practice (nonsignificant Type * Period interaction, *F*(2, 70) = 1.54, *p* = .221, η_*p*_^2^ = .042; see Fig. 4A, B). Meanwhile, as shown by the significant Period main effect, *F*(2, 70) = 13.24, ε = .715, *p* < .001, η_*p*_^2^ = .274, the overall CNV amplitude gradually increased as the task progressed (period_1_: −0.44 μV, period_2_: −0.84 μV, period_3_: −1.05 μV; all pairwise differences were significant at *p* ≤ .014; see Fig. 2B and Fig. 4A).

#### 3.2.2 P3 peak

The Type by Period ANOVA on the P3 peak showed a significant main effect of Type, *F*(1, 35) = 11.03, *p* = .002, η_*p*_^2^ = .240, indicating that the P3 peak amplitude was lower for random low-probability triplets than for pattern high-probability ones (3.40 μV vs. 3.54 μV; see Fig. 2C and Fig. 3B, C). This difference did not change with practice (nonsignificant Type * Period interaction, *F*(2, 70) = 1.43, *p* = .246, η_*p*_^2^ = .039; see Fig 4A, B). The significant Period main effect, *F*(2, 70) = 4.64, ε = .708, *p* = .024, η_*p*_^2^ = .117, showed that the overall P3 peak amplitude decreased as the task progressed (period_1_: 3.70 μV, period_2_: 3.47 μV, period_3_: 3.25 μV; period_1_ > period_3,_ *p* = .021, period_2_ > period_3_, *p* = .045; see Fig. 2C and Fig. 4A). In terms of the Type main effect, comparable results were obtained when the response-locked P3 peak was analyzed (see Supplementary Material).

#### 3.2.3 Late P3

Similar results emerged for the late P3 from the Type by Period ANOVA. The mean amplitude of the late P3 was lower for random low-probability triplets than for pattern high-probability ones (significant main effect of Type, *F*(1, 35) = 13.51, *p* = .001, η_*p*_^2^ = .278; 2.07 μV vs. 2.23 μV; see Fig. 2D and Fig. 3B, C), and, irrespective of the triplet type, the overall mean amplitude of the late P3 decreased with practice (significant main effect of Period, *F*(2, 70) = 11.26, ε = .804, *p* < .001, η_*p*_^2^ = .243; period_1_: 2.49 μV, period_2_: 2.05 μV, period_3_: 1.90 μV; period_1_ > period_2,_ *p* = .002, period_1_ > period_3_, *p* < .001; see Fig. 2D and Fig. 4A). The Type * Period interaction was nonsignificant, *F*(2, 70) = 0.06, *p* = .942, η_*p*_^2^ = .002 (see Fig. 4A, B). In terms of the Type main effect, comparable results were obtained when the late phase of the response-locked P3 was analyzed (see Supplementary Material).

### 3.3 Correlations between the behavioral and ERP measures

Correlational analyses were performed to test (1) the potential speed-accuracy trade-off in task performance and (2) whether behavioral measures were related to the ERPs. Therefore, associations (Pearson correlations) between (1) RTs and accuracy and (2) between each of these behavioral measures and the CNV, P3 peak, and late P3 amplitudes were investigated. These analyses were run for each triplet type separately in the corresponding time periods (e.g., RTs to random low-probability triplets in period_2_ with CNV amplitude for random low-probability triplets in period_2_) as well as for the entire acquisition phase (e.g., mean accuracy for pattern high-probability triplets with the mean amplitude of the CNV for pattern high-probability triplets).

None of the correlations were significant between RTs and accuracy (all |*r*|s ≤ .150, *p*s ≥ .382), indicating the lack of speed-accuracy trade-off. Similarly, no significant correlations were found between the behavioral measures and the CNV amplitude (all |*r*|s ≤ .166, *p*s ≥ .334). In contrast, significant negative correlations consistently emerged between RTs and the P3 peak, indicating that faster responses were associated with larger P3 peak amplitudes (pattern high-probability triplets: period_1_ *r*(34) = −.449, *p* = .006; period_2_ *r*(34) = −.516, *p* = .001; period_3_ *r*(34) = −.456, *p* = .005; all periods: *r*(34) = −.485, *p* = .003; random low-probability triplets: period_1_ *r*(34) = −.455, *p* = .005; period_2_ *r*(34) = −.448, *p* = .006; period_3_ *r*(34) = −.405, *p* = .014; all periods: *r*(34) = −.443, *p* = .007). However, this was not observed in the case of accuracy (all |*r*|s ≤ .304, *p*s ≥ .071; but see the response-locked averages in the Supplementary Material). In terms of the response-locked P3 peak, a similar pattern of associations emerged with the RTs, albeit the correlations were weaker and less consistent (the correlations were significant only in the first two periods for pattern high-probability triplets and in the first period for random low-probability triplets, see Supplementary Material). No significant correlations were found between the behavioral measures and the amplitude of the late P3 (neither in the cue-locked nor in the response-locked averages; all |*r*|s ≤ .216, *p*s ≥ .205).

## 4. Discussion

### 4.1 Summary of results

This study provided ERP evidence for the operation of implicit anticipatory processes in the acquisition of probabilistic regularities. The CNV developed during the presentation of the cue until target onset, and its amplitude was larger before unpredictable than before predictable targets. Importantly, the cue was not predictive of the target. The ERP difference reflecting sensitivity to the underlying regularities was continuously present with the same magnitude in the time window of target processing, indicated by the amplitudes of the P3 peak and the late P3 being lower for unpredictable than for predictable targets. As expected, anticipatory processing was shown not only by the CNV but also by the RTs that were faster to predictable targets. Based on these results, we propose that internal models have been implicitly formed on the probabilistic regularities of the stimulus stream. These internal models could have guided a series of anticipatory and preparatory processes that eventually facilitated stimulus processing.

### 4.2 Interpretation of behavioral results

Behavioral evidence accumulated so far indicates that participants respond overall faster or increasingly faster to high-probability triplets than to low-probability ones in the ASRT task (e.g., Janacsek et al., 2015; Kóbor et al., 2017; Nemeth et al., 2010; Nemeth, Janacsek, Polner, et al., 2013; Szegedi-Hallgató et al., 2019; Takács et al., 2017; Tóth et al., 2017). Faster responses specifically to pattern high-probability triplets than to random low-probability ones were also observed in task versions without target-preceding cues (Horváth et al., in press; Kóbor et al., 2019; Kóbor et al., 2018; Nemeth, Janacsek, & Fiser, 2013; Szegedi-Hallgató et al., 2019). The magnitude of the present RT effect was small (2 ms) relative to other studies using the ASRT task. This can be explained by the combination of the following factors: the present experiment used a slow-paced task version with fixed ITI, acquisition occurred in an implicit and incidental manner, and the target-preceding cue not only lengthened the ITI but could have also influenced acquisition in a currently unknown manner. Therefore, irrespective of the reliability of the RT effect, it is expected to be smaller or larger based on the combination of these factors as described below.

Larger RT effects (ca. 10-15 ms) have been generally observed in self-paced ASRT tasks with fast stimulus presentation rate, in which the acquisition of probabilities has occurred in an incidental manner (Kóbor et al., 2017; Nemeth et al., 2010; Stark-Inbar, Raza, Taylor, & Ivry, 2017). These RT effects could further increase in ASRT tasks where the acquisition of probabilities is intentional (Nemeth, Janacsek, & Fiser, 2013; Simor et al., 2019; Tóth-Fáber, Janacsek, Szőllősi, Kéri, & Németh, 2020). Meanwhile, in fixed-paced tasks (Kóbor et al., 2020; Tóth et al., 2017) and in those with slow stimulus presentation rate (i.e., long ITI, see Kiss, Nemeth, & Janacsek, 2019; Kóbor et al., 2019), RT effects have been found to be reduced (but see Horváth, Török, Pesthy, Nemeth, & Janacsek, 2020). The present task was fixed-paced, the length of the ITI was longer than previously yielding a slow rate of stimulus presentation, and acquisition occurred incidentally. These characteristics could have altogether reduced the magnitude of the RT effect. Moreover, albeit a target-preceding cue stimulus in the ASRT task has not been applied so far, one should also take into account the impact of the cue when explaining effect size differences between the studies.

Accordingly, we assume that since the stimulus presentation rate was slower than in other studies, the incidental binding of successive trials (i.e., triplet elements) could have been weaker. The insertion of an irrelevant cue stimulus into the trial structure could have further distorted or at least interfered with the binding process. This weaker binding possibly altered the extraction of the triplet-level probability structure, eventually yielding reduced measures of implicit acquisition. Alternatively, slow stimulus presentation rate and the insertion of the cue could have decreased the facilitatory effect of predictable stimulus combinations (i.e., high-probability triplets). Particularly, when participants responded to the current target, the representations of previous stimuli might have been inactivated, resulting in weaker response facilitation (for more details on this interpretation, see Kiss et al., 2019). As a result of weaker response facilitation, although participants possibly acquired the probability structure to some degree, the momentary expression of this acquired knowledge was hindered with slow stimulus presentation rate (Kiss et al., 2019; Willingham, Greenberg, & Thomas, 1997). Based on the behavioral pilot study of the current experiment, the latter interpretation seems the most likely. Particularly, when stimulus presentation was changed from the original slower rate (1800 ms ITI) to a faster one (900 ms ITI) in the pilot study, larger RT effects were observed.

Nevertheless, as RTs differed between pattern high-probability and random low-probability triplets and this difference stabilized by the end of the task, the conclusions drawn from the behavioral data are still in line with earlier behavioral (cf. Juhasz et al., 2019; Kóbor et al., 2020; Szegedi-Hallgató et al., 2019; Török, Janacsek, Nagy, Orbán, & Nemeth, 2017) and ERP findings (Kóbor et al., 2019; Kóbor et al., 2018). The latter studies together with the present one suggest that the sensitivity to multiple probabilistic regularities has possibly been grounded in the implicit extraction of the triplet-level probability structure.

### 4.3 Interpretation of ERP results

#### 4.3.1 CNV

As in the present study, larger CNV for unpredictable vs. predictable visual stimuli was found previously during the implicit acquisition of probabilistic relations (Rose et al., 2001). Meanwhile, studies involving more easily learnable sequences of auditory stimuli (Jongsma et al., 2006; Stadler et al., 2006) and those that contrasted predictive auditory contexts with random ones in explicit experimental settings (Fogelson et al., 2009) showed the opposite. However, in those studies, repeating predictor/standard–target sequences were used. These sequences differ from the one we applied in the present implicit ASRT task version: Here, a stream of perceptually identical, task-relevant targets that all require keypresses create the ongoing stimulus context, and the exact directions of the targets are determined only by the underlying regularity. In our view, the processes reflected by the CNV in our task are related to the pre-allocation of attentional resources and to response preparation in an implicit and nonconscious manner. We base this interpretation on the assumption that the random low-probability, unpredictable stimuli would be more complex to-be-responded targets than the pattern high-probability ones, as already observed at the behavioral level. The potential functional role of the CNV in the current setting is further elaborated below.

The observed centroparietal distribution of the CNV difference between unpredictable and predictable targets might originate from the characteristics of the current task and the neurocognitive processes underlying performance. As in our study, the late CNV is more centrally distributed and involves a posterior cortical network in other perceptual-motor tasks (e.g., Berchicci et al., 2016; Brunia & Damen, 1988; De Kleine & Van der Lubbe, 2011; Gómez et al., 2003; Leynes et al., 1998). Furthermore, it has been suggested that the late CNV reflects two aspects of the parieto-frontal motor system: the maintenance of the stimulus-response (S–R) links and the activation of the hand-motor area (Verleger et al., 2000). Therefore, the observed CNV difference in the present study might indicate the binding of stimulus information with the related response to help the selection between different motor programs. However, beyond response preparation, implicit and nonconscious predictions were likely generated about the direction of the upcoming target (cf. Kóbor et al., 2020; Kóbor et al., 2019). Consequently, it is conceivable that the presentation of the target arrow per se served as a feedback about the “correctness” of these predictions (i.e., whether the targets were anticipated outcomes or not). Thus, the late CNV could have also consisted of the stimulus-preceding negativity (SPN) reflecting anticipatory attention to the perceptual informational delivered by the target, which might additionally contribute to the more posterior scalp distribution of the difference (Brunia, Hackley, van Boxtel, Kotani, & Ohgami, 2011; Hackley et al., 2014; van Boxtel & Brunia, 1994).

We intended to measure the genuine predictability of the targets because of acquiring the probability structure of the task. To this end, a simple fixation cross was used as the cue, which is somewhat unusual in cued RT tasks (but see Boehm et al., 2014; Pauletti et al., 2014). The fixation cross did not contain probability or direction information about the upcoming target. Since no information was delivered about which S–R links should be selected before target presentation, a generic pre-activation of the sensory-motor areas might have been needed for fast and accurate responding (Pauletti et al., 2014). Similarly, sustained activity in the frontoparietal network mediating response selection (cf. Schumacher, Hendricks, & D’Esposito, 2005; Verleger et al., 2000) might have also been involved in task solving. Supporting this notion, the amplitude of the CNV was not only modulated as a function of target probability but it also increased as the task progressed in time, irrespective of probability information. In our earlier study, when the pattern high-probability triplets were perceptually distinguishable from the random ones (i.e., marked with different colors) and no other cues were used, proficiency in the task and response automatization for the former triplets were evidenced by the decreasing amplitude of different ERP components (N2, P3, Ne, Horváth et al., in press; Kóbor et al., 2018). In contrast, since target probability was not explicitly denoted in the present experiment, and the task was perceived as increasingly exhausting according to verbal reports, the probability-non-specific CNV amplitude enhancement likely indicates the growing effort participants needed to maintain task performance. Sustained effort needed for stimulus processing and responding is also reflected in the RTs. Although RTs became faster due to practice at the descriptive level, general skill improvements were not statistically significant over task periods. (Note that general skill improvements are independent of the observed RT difference between high- and low-probability triplets.) According to the correlational results, no speed-accuracy trade-off characterized task performance, and it is unlikely that the CNV amplitude enhancement was associated with lower response caution and/or increased response speed (cf. Boehm et al., 2014).

#### 4.3.2 P3

The differential ERP effect for predictable vs. unpredictable stimuli persisted in the time window of the P3 component with similar magnitude. Particularly, the enhanced ERP amplitudes before unpredictable target stimuli appeared as decreased amplitudes after target onset (cf. Killikelly & Szűcs, 2013). The P3 amplitude has long been suggested to scale the probability of the stimulus (Donchin, 1981; Donchin & Coles, 1988; Mars et al., 2008) and to be modulated by the acquisition of predictive relations embedded in the stimulus sequence (e.g., Batterink, Reber, Neville, & Paller, 2015; Eimer, Goschke, Schlaghecken, & Stürmer, 1996; Jost, Conway, Purdy, Walk, & Hendricks, 2015; Rüsseler, Münte, & Wiswede, 2018). Meanwhile, current theoretical accounts of this component suggest its more intricate functional relevance in decision making and in linking stimulus evaluation to response selection (Kelly & O’Connell, 2015; Twomey, Murphy, Kelly, & O’Connell, 2015; Verleger, Jaśkowski, & Wascher, 2005; Verleger & Śmigasiewicz, 2016).

Particularly, the P3 component could be considered as reflecting the process of mapping a task-relevant stimulus onto an appropriate response (Berchicci et al., 2016; Folstein & van Petten, 2011; Stock, Steenbergen, Colzato, & Beste, 2016; Verleger et al., 2005). Analysis of the response-locked averages strengthens this notion by showing the differentiation of predictable vs. unpredictable stimuli also in the response-locked P3 component (see Supplementary Material and note that at the descriptive level, this effect is slightly weaker in the response-locked than in the cue-locked averages, especially in the case of the P3 peak). The relation between the P3 component and motor responses were further supported by the negative correlations emerging between RTs and the P3 peak (in both the cue-locked and response-locked averages), indicating faster responses associated with larger P3 peaks, irrespective of the predictability of the target.

In terms of the theoretical accounts of the P3, the concept of S–R link might help explaining the P3 findings. According to this concept, S–R links established with practice during the task are reactivated for initiating the correct response, and the P3 reflects the amount of this reactivation process (e.g., Verleger, Hamann, Asanowicz, & Śmigasiewicz, 2015; Verleger et al., 2017). The implicit acquisition of the probability structure could change the S–R links related to unpredictable stimuli that might require stronger reactivation, yielding larger P3 amplitudes (cf. Kóbor et al., 2019). However, due to the cued design, some of the stimulus processing might occur in an anticipatory manner. It is possible that the stronger response preparation before the unpredictable stimuli facilitates target processing that uses less attentional resources (Polich, 2007; Polich & Criado, 2006) and operates at a lower decision threshold (Kelly & O’Connell, 2015), eventually yielding *decreased* P3 amplitudes. Decreased P3 amplitudes have been found to reflect decision uncertainty and that the resources for stimulus processing are needed elsewhere during effortful processing (Beauducel, Brocke, & Leue, 2006; Johnson, 1986; Kok, 2001), which could also explain the increased CNV and decreased P3 for unpredictable stimuli. At the same time, the facilitated “pre-processing” is not reflected in the RTs, as those are still slower for unpredictable stimuli. The implicit, nonconscious aspect of the task, however, might explain this matter, since “implicit preparation” might be enough for responding correctly but not for responding as fast as to predictable targets. Nevertheless, this finding is also in line with the assumption that movement initiation and movement preparation could be independent processes (Haith, Pakpoor, & Krakauer, 2016).

#### 4.3.3 Mutual properties of the CNV and P3 effects

As delineated above, the CNV and the P3 seem to be interrelated. Accordingly, the observed ERP variation in the P3 time window can be regarded as a “CNV return” to baseline and its “overshoot” into the positive direction (Verleger et al., 2013). To further support this concept, not only the ERP effect related to predictability persisted after target onset but also the time-on-task effect, i.e., the overall CNV enhancement, which appeared as decreasing P3 amplitudes as the task progressed. Although the response-locked P3 amplitudes did not change in time, the CNV also appeared to be time-invariant in those averages (see Supplementary Figs. S1-S2), additionally indicating associations between the two components. Moreover, although the CNV looks mostly unaffected by ERP locking (Berchicci et al., 2016), response-locked waveforms imply that the CNV is more likely related to the target stimulus than to the response: It develops until target onset and returns to baseline before response onset. This observation does not preclude that the CNV amplitude modulation possibly reflects the activation and maintenance of the S–R link relevant to the given target (Verleger et al., 2000).

Similar to the RT effects, the magnitudes of the observed CNV and P3 effects were also small (0.14–0.16 μV) relative to other independent studies on the acquisition of probabilistic regularities (e.g., Batterink et al., 2015; Emerson et al., 2014; Rüsseler et al., 2018). This phenomenon might be explained by those characteristics of the ASRT task and underlying processes that we described in relation to the RT effects (see also the smaller ERP effects in ASRT tasks with slower stimulus presentation rate, Kóbor et al., 2019; Kóbor et al., 2018). Indeed, as acquisition occurred incidentally in the present experiment, smaller ERP effects were expected. Earlier studies using variants of the serial reaction time task combined with features of the oddball paradigm showed smaller, altered, or lacking ERP effects for incidental as compared with intentional acquisition (e.g., Eimer et al., 1996; Ferdinand, Mecklinger, & Kray, 2008; Rüsseler, Hennighausen, Münte, & Rösler, 2003; Rüsseler & Rösler, 2000). In line with this notion, the magnitude of the present P3 differences is comparable to those observed in our previous study where acquisition was incidental (Kóbor et al., 2019). Although paradigms structurally different from the ASRT task might promote the occurrence of larger CNV and P3 effects (cf. Batterink et al., 2015; Emerson et al., 2014; Jost et al., 2015; Rose et al., 2001), the significance of our ERP findings is that they indicate the implicit anticipation of predictable and unpredictable events in an unsupervised statistical learning environment.

### 4.4 Conclusions

To conclude, the behavioral and ERP markers of implicit anticipation differed between predictable and unpredictable events arranged in a sequence of visual stimuli. Although the cue signals were unspecific to the predictability of these target events, lengthened RTs together with enhanced CNV and attenuated P3 amplitudes were recorded for unpredictable as compared with predictable events. To explain these ERP amplitude modulations, we propose that, during the preparatory phase of processing, more resources were needed to maintain the stimulus-response contingencies appropriate for the unpredictable events. This pattern of results possibly emerged due to the implicit and incidental acquisition of the probabilistic regularities through the formation of internal models, which is fundamental not only in perceptual and response selection processes but also in learning and memory. In line with this interpretation, our results further suggest that the CNV, by reflecting the active operation of these internal models, could be considered as a neurocognitive marker of uncertainty. This ERP component would be useful in investigating the acquisition of probabilistic regularities in clinical and non-clinical samples.

## Acknowledgement

This research was supported by the National Brain Research Program (project 2017-1.2.1-NKP-2017-00002, PI: D. N.); the Hungarian Scientific Research Fund (OTKA FK 124412, PI: A. K., OTKA PD 124148, PI: K. J., OTKA K 128016, PI: D. N.); the IDEXLYON Fellowship of the University of Lyon as part of the Programme Investissements d’Avenir (ANR-16-IDEX-0005 to D. N.); the János Bolyai Research Scholarship of the Hungarian Academy of Sciences (to A. K. and K. J.); and the ÚNKP-19-3 New National Excellence Program of the Ministry for Innovation and Technology (to K. H.). The authors thank the help of Dorottya Denke in data acquisition. They also thank Márk Molnár and Zsófia Anna Gaál for their pieces of advice on the design, ERP analysis, and initial results.

## Disclosure of interest

The authors report no conflicts of interest.

## Notes

### Competing Interest Statement

The authors have declared no competing interest.

### Summary of Updates

Added the supplementary material.

## References

Anatürk, M., & Jentzsch, I. (2015). The effects of musical training on movement pre-programming and re-programming abilities: An event-related potential investigation. Biological Psychology, 106, 39–49. doi:10.1016/j.biopsycho.2015.01.014

Armstrong, B. C., Frost, R., & Christiansen, M. H. (2017). The long road of statistical learning research: Past, present and future. Philosophical Transactions of the Royal Society B: Biological Sciences, 372(1711). doi:10.1098/rstb.2016.0047

Aslin, R. N. (2017). Statistical learning: A powerful mechanism that operates by mere exposure. Wiley Interdisciplinary Reviews: Cognitive Science, 8(1-2), e1373. doi:10.1002/wcs.1373

Batterink, L. J., Reber, P. J., Neville, H. J., & Paller, K. A. (2015). Implicit and explicit contributions to statistical learning. Journal of Memory and Language, 83, 62–78. doi:10.1016/j.jml.2015.04.004

Beauducel, A., Brocke, B., & Leue, A. (2006). Energetical bases of extraversion: Effort, arousal, EEG, and performance. International Journal of Psychophysiology, 62(2), 212–223. doi:10.1016/j.ijpsycho.2005.12.001

Berchicci, M., Spinelli, D., & Di Russo, F. (2016). New insights into old waves. Matching stimulus- and response-locked ERPs on the same time-window. Biological Psychology, 117, 202–215. doi:10.1016/j.biopsycho.2016.04.007

Boehm, U., van Maanen, L., Forstmann, B., & van Rijn, H. (2014). Trial-by-trial fluctuations in CNV amplitude reflect anticipatory adjustment of response caution. Neuroimage, 96, 95–105. doi:10.1016/j.neuroimage.2014.03.063

Brunia, C. H. M. (2003). Cnv and spn: Indices of anticipatory behavior. In M. Jahanshahi & M. Hallett (Eds.), The Bereitschaftspotential: Movement-related cortical potentials (pp. 207–227). Boston, MA: Springer US.

Brunia, C. H. M., & Damen, E. J. P. (1988). Distribution of slow brain potentials related to motor preparation and stimulus anticipation in a time estimation task. Electroencephalography and Clinical Neurophysiology, 69(3), 234–243. doi:10.1016/0013-4694(88)90132-0

Brunia, C. H. M., Hackley, S. A., van Boxtel, G. J., Kotani, Y., & Ohgami, Y. (2011). Waiting to perceive: Reward or punishment? Clinical Neurophysiology, 122(5), 858–868. doi:10.1016/j.clinph.2010.12.039

Conway, C. M. (2020). How does the brain learn environmental structure? Ten core principles for understanding the neurocognitive mechanisms of statistical learning. Neuroscience & Biobehavioral Reviews. doi:10.1016/j.neubiorev.2020.01.032

Cui, R. Q., Egkher, A., Huter, D., Lang, W., Lindinger, G., & Deecke, L. (2000). High resolution spatiotemporal analysis of the contingent negative variation in simple or complex motor tasks and a non-motor task. Clinical Neurophysiology, 111(10), 1847–1859. doi:10.1016/s1388-2457(00)00388-6

Daltrozzo, J., & Conway, C. M. (2014). Neurocognitive mechanisms of statistical-sequential learning: What do event-related potentials tell us? Frontiers in Human Neuroscience, 8(437). doi:10.3389/fnhum.2014.00437

Daw, N. D., Gershman, S. J., Seymour, B., Dayan, P., & Dolan, R. J. (2011). Model-based influences on humans’ choices and striatal prediction errors. Neuron, 69(6), 1204–1215. doi:10.1016/j.neuron.2011.02.027

De Kleine, E., & Van der Lubbe, R. H. (2011). Decreased load on general motor preparation and visual-working memory while preparing familiar as compared to unfamiliar movement sequences. Brain and Cognition, 75(2), 126–134. doi:10.1016/j.bandc.2010.10.013

Delorme, A., Sejnowski, T., & Makeig, S. (2007). Enhanced detection of artifacts in EEG data using higher-order statistics and independent component analysis. Neuroimage, 34(4), 1443–1449. doi:10.1016/j.neuroimage.2006.11.004

Di Russo, F., Berchicci, M., Bozzacchi, C., Perri, R. L., Pitzalis, S., & Spinelli, D. (2017). Beyond the “Bereitschaftspotential”: Action preparation behind cognitive functions. Neuroscience & Biobehavioral Reviews, 78, 57–81. doi:10.1016/j.neubiorev.2017.04.019

Donchin, E. (1981). Surprise!… surprise? Psychophysiology, 18(5), 493–513. doi:doi:10.1111/j.1469-8986.1981.tb01815.x

Donchin, E., & Coles, M. G. H. (1988). Is the P300 component a manifestation of context updating? Behavioral and Brain Sciences, 11(3), 357–374. doi:10.1017/S0140525X00058027

Dragovic, M. (2004a). Categorization and validation of handedness using latent class analysis. Acta Neuropsychiatrica, 16(4), 212–218. doi:10.1111/j.0924-2708.2004.00087.x

Dragovic, M. (2004b). Towards an improved measure of the Edinburgh Handedness Inventory: A one-factor congeneric measurement model using confirmatory factor analysis. Laterality: Asymmetries of Body, Brain and Cognition, 9(4), 411–419. doi:10.1080/13576500342000248

Eimer, M., Goschke, T., Schlaghecken, F., & Stürmer, B. (1996). Explicit and implicit learning of event sequences: Evidence from event-related brain potentials. Journal of Experimental Psychology. Learning, Memory, and Cognition, 22(4), 970–987. doi:10.1037/0278-7393.22.4.970

Emerson, S., Daltrozzo, J., & Conway, C. (2014). The effect of music experience on auditory sequential learning: An erp study. Proceedings of the Annual Meeting of the Cognitive Science Society, 36, 2157–2162. Retrieved from https://escholarship.org/uc/item/28t877px

Fan, J., Kolster, R., Ghajar, J., Suh, M., Knight, R. T., Sarkar, R., & McCandliss, B. D. (2007). Response anticipation and response conflict: An event-related potential and functional magnetic resonance imaging study. The Journal of Neuroscience, 27(9), 2272–2282. doi:10.1523/jneurosci.3470-06.2007

Ferdinand, N. K., Mecklinger, A., & Kray, J. (2008). Error and deviance processing in implicit and explicit sequence learning. Journal of Cognitive Neuroscience, 20(4), 629–642. doi:10.1162/jocn.2008.20046

Fogelson, N., Wang, X., Lewis, J. B., Kishiyama, M. M., Ding, M., & Knight, R. T. (2009). Multimodal effects of local context on target detection: Evidence from P3b. Journal of Cognitive Neuroscience, 21(9), 1680–1692. doi:10.1162/jocn.2009.21071

Folstein, J. R., & van Petten, C. (2011). After the P3: Late executive processes in stimulus categorization. Psychophysiology, 48(6), 825–841. doi:10.1111/j.1469-8986.2010.01146.x

Friston, K. (2005). A theory of cortical responses. Philosophical transactions of the Royal Society of London. Series B, Biological sciences, 360(1456), 815–836. doi:10.1098/rstb.2005.1622

Friston, K. (2010). The free-energy principle: A unified brain theory? Nature Reviews Neuroscience, 11(2), 127–138. doi:10.1038/nrn2787

Friston, K. J., Stephan, K. E., Montague, R., & Dolan, R. J. (2014). Computational psychiatry: The brain as a phantastic organ. The Lancet Psychiatry, 1(2), 148–158. doi:10.1016/S2215-0366(14)70275-5

Frost, R., Armstrong, B. C., & Christiansen, M. H. (2019). Statistical learning research: A critical review and possible new directions. Psychological Bulletin, 145(12), 1128–1153. doi:10.1037/bul0000210

Gómez, C. M., & Flores, A. (2011). A neurophysiological evaluation of a cognitive cycle in humans. Neuroscience and Biobehavioral Reviews, 35(3), 452–461. doi:10.1016/j.neubiorev.2010.05.005

Gómez, C. M., Marco, J., & Grau, C. (2003). Preparatory visuo-motor cortical network of the contingent negative variation estimated by current density. Neuroimage, 20(1), 216–224. doi:10.1016/S1053-8119(03)00295-7

Greenhouse, S., & Geisser, S. (1959). On methods in the analysis of profile data. Psychometrika, 24(2), 95–112. doi:10.1007/bf02289823

Griffiths, T. L., Kemp, C., & Tenenbaum, J. B. (2008). Bayesian models of cognition. In R. Sun (Ed.), Cambridge handbook of computational cognitive modeling: Cambridge University Press.

Hackley, S. A., Valle-Inclán, F., Masaki, H., & Hebert, K. (2014). Stimulus-preceding negativity (SPN) and attention to rewards. In G. R. Mangun (Ed.), Cognitive electrophysiology of attention: Signals of the mind (pp. 216–225). San Diego: Academic Press.

Haith, A. M., Pakpoor, J., & Krakauer, J. W. (2016). Independence of movement preparation and movement initiation. The Journal of Neuroscience, 36(10), 3007. doi:10.1523/JNEUROSCI.3245-15.2016

Hallgató, E., Győri-Dani, D., Pekár, J., Janacsek, K., & Nemeth, D. (2013). The differential consolidation of perceptual and motor learning in skill acquisition. Cortex, 49(4), 1073–1081. doi:10.1016/j.cortex.2012.01.002

Hommel, B. (2009). Action control according to TEC (theory of event coding). Psychological Research, 73(4), 512–526. doi:10.1007/s00426-009-0234-2

Horváth, K., Kardos, Z., Takács, Á., Csépe, V., Nemeth, D., Janacsek, K., & Kóbor, A. (in press). Error processing during the online retrieval of probabilistic sequence knowledge. Journal of Psychophysiology. doi:10.1027/0269-8803/a000262

Horváth, K., Török, C., Pesthy, O., Nemeth, D., & Janacsek, K. (2020). Divided attention does not affect the acquisition and consolidation of transitional probabilities. Scientific Reports, 10(1), 22450. doi:10.1038/s41598-020-79232-y

Howard, D. V., Howard, J. H., Jr., Japikse, K., DiYanni, C., Thompson, A., & Somberg, R. (2004). Implicit sequence learning: Effects of level of structure, adult age, and extended practice. Psychology and Aging, 19(1), 79–92. doi:10.1037/0882-7974.19.1.79

Howard, J. H., Jr., & Howard, D. V. (1997). Age differences in implicit learning of higher order dependencies in serial patterns. Psychology and Aging, 12(4), 634–656. doi:10.1037/0882-7974.12.4.634

Janacsek, K., Ambrus, G. G., Paulus, W., Antal, A., & Nemeth, D. (2015). Right hemisphere advantage in statistical learning: Evidence from a probabilistic sequence learning task. Brain Stimulation, 8(2), 277–282. doi:10.1016/j.brs.2014.11.008

Jentzsch, I., Leuthold, H., & Richard ridderinkhof, K. (2004). Beneficial effects of ambiguous precues: Parallel motor preparation or reduced premotoric processing time? Psychophysiology, 41(2), 231–244. doi:10.1111/j.1469-8986.2004.00155.x

Johnson, R. (1986). A triarchic model of P300 amplitude. Psychophysiology, 23(4), 367–384. doi:10.1111/j.1469-8986.1986.tb00649.x

Jongsma, M. L. A., Eichele, T., Van Rijn, C. M., Coenen, A. M. L., Hugdahl, K., Nordby, H., & Quiroga, R. Q. (2006). Tracking pattern learning with single-trial event-related potentials. Clinical Neurophysiology, 117(9), 1957–1973. doi:10.1016/j.clinph.2006.05.012

Jost, E., Conway, C. M., Purdy, J. D., Walk, A. M., & Hendricks, M. A. (2015). Exploring the neurodevelopment of visual statistical learning using event-related brain potentials. Brain Research, 1597, 95–107. doi:10.1016/j.brainres.2014.10.017

Juhasz, D., Nemeth, D., & Janacsek, K. (2019). Is there more room to improve? The lifespan trajectory of procedural learning and its relationship to the between- and within-group differences in average response times. PLoS ONE, 14(7), e0215116. doi:10.1371/journal.pone.0215116

Kelly, S. P., & O’Connell, R. G. (2015). The neural processes underlying perceptual decision making in humans: Recent progress and future directions. Journal of Physiology-Paris, 109(1), 27–37. doi:10.1016/j.jphysparis.2014.08.003

Killikelly, C., & Szűcs, D. (2013). Delayed development of proactive response preparation in adolescents: ERP and EMG evidence. Developmental Cognitive Neuroscience, 3, 33–43. doi:10.1016/j.dcn.2012.08.002

Kiss, M., Nemeth, D., & Janacsek, K. (2019). Stimulus presentation rates affect performance but not the acquired knowledge – evidence from procedural learning. bioRxiv, 650598. doi:10.1101/650598

Kóbor, A., Horváth, K., Kardos, Z., Nemeth, D., & Janacsek, K. (2020). Perceiving structure in unstructured stimuli: Implicitly acquired prior knowledge impacts the processing of unpredictable transitional probabilities. Cognition, 205, 104413. doi:10.1016/j.cognition.2020.104413

Kóbor, A., Horváth, K., Kardos, Z., Takács, Á., Janacsek, K., Csépe, V., & Nemeth, D. (2019). Tracking the implicit acquisition of nonadjacent transitional probabilities by ERPs. Memory & Cognition, 47(8), 1546–1566. doi:10.3758/s13421-019-00949-x

Kóbor, A., Janacsek, K., Takács, Á., & Nemeth, D. (2017). Statistical learning leads to persistent memory: Evidence for one-year consolidation. Scientific Reports, 7(1), 760. doi:10.1038/s41598-017-00807-3

Kóbor, A., Takács, Á., Kardos, Z., Janacsek, K., Horváth, K., Csépe, V., & Nemeth, D. (2018). Erps differentiate the sensitivity to statistical probabilities and the learning of sequential structures during procedural learning. Biological Psychology, 135, 180–193. doi:10.1016/j.biopsycho.2018.04.001

Koelsch, S., Busch, T., Jentschke, S., & Rohrmeier, M. (2016). Under the hood of statistical learning: A statistical mmn reflects the magnitude of transitional probabilities in auditory sequences. Scientific Reports, 6, 19741. doi:10.1038/srep19741

Kok, A. (2001). On the utility of P3 amplitude as a measure of processing capacity. Psychophysiology, 38(3), 557–577. doi:10.1017/S0048577201990559

Kononowicz, T. W., & Penney, T. B. (2016). The contingent negative variation (CNV):Timing isn’t everything. Current Opinion in Behavioral Sciences, 8, 231–237. doi:10.1016/j.cobeha.2016.02.022

Leuthold, H., & Jentzsch, I. (2002). Spatiotemporal source localisation reveals involvement of medial premotor areas in movement reprogramming. Experimental Brain Research, 144(2), 178–188. doi:10.1007/s00221-002-1043-7

Leuthold, H., Sommer, W., & Ulrich, R. (2004). Preparing for action: Inferences from CNV and LRP. Journal of Psychophysiology, 18(2-3), 77–88. doi:10.1027/0269-8803.18.23.77

Leynes, P. A., Allen, J. D., & Marsh, R. L. (1998). Topographic differences in CNV amplitude reflect different preparatory processes. International Journal of Psychophysiology, 31(1), 33–44. doi:10.1016/S0167-8760(98)00032-4

Loveless, N. E., & Sanford, A. J. (1974). Slow potential correlates of preparatory set. Biological Psychology, 1(4), 303–314. doi:10.1016/0301-0511(74)90005-2

Macar, F., & Vidal, F. (2004). Event-related potentials as indices of time processing: A review. Journal of Psychophysiology, 18(2/3), 89–104. doi:10.1027/0269-8803.18.23.89

Maheu, M., Dehaene, S., & Meyniel, F. (2019). Brain signatures of a multiscale process of sequence learning in humans. eLife, 8, e41541. doi:10.7554/eLife.41541

Mars, R. B., Debener, S., Gladwin, T. E., Harrison, L. M., Haggard, P., Rothwell, J. C., & Bestmann, S. (2008). Trial-by-trial fluctuations in the event-related electroencephalogram reflect dynamic changes in the degree of surprise. The Journal of Neuroscience, 28(47), 12539–12545. doi:10.1523/jneurosci.2925-08.2008

Medimorec, S., Milin, P., & Divjak, D. (2019). Working memory affects anticipatory behavior during implicit pattern learning. Psychological Research. doi:10.1007/s00426-019-01251-w

Meyniel, F., Maheu, M., & Dehaene, S. (2016). Human inferences about sequences: A minimal transition probability model. PLoS Computational Biology, 12(12), e1005260–e1005260. doi:10.1371/journal.pcbi.1005260

Molnár, M., Csuhaj, R., Gaál, Z. A., Czigler, B., Ulbert, I., Boha, R., & Kondákor, I. (2008). Spectral characteristics and linear-nonlinear synchronization changes of different EEG frequency bands during the CNV. Psychophysiology, 45(3), 412–419. doi:10.1111/j.1469-8986.2008.00648.x

Nemeth, D., & Janacsek, K. (2011). The dynamics of implicit skill consolidation in young and elderly adults. The Journals of Gerontology. Series B, Psychological Sciences and Social Sciences, 66(1), 15–22. doi:10.1093/geronb/gbq063

Nemeth, D., Janacsek, K., & Fiser, J. (2013). Age-dependent and coordinated shift in performance between implicit and explicit skill learning. Frontiers in Computational Neuroscience, 7, 147. doi:10.3389/fncom.2013.00147

Nemeth, D., Janacsek, K., Londe, Z., Ullman, M. T., Howard, D. V., & Howard, J. H., Jr. (2010). Sleep has no critical role in implicit motor sequence learning in young and old adults. Experimental Brain Research, 201(2), 351–358. doi:10.1007/s00221-009-2024-x

Nemeth, D., Janacsek, K., Polner, B., & Kovacs, Z. A. (2013). Boosting human learning by hypnosis. Cerebral Cortex, 23(4), 801–805. doi:10.1093/cercor/bhs068

Oldfield, R. C. (1971). The assessment and analysis of handedness: The Edinburgh Inventory. Neuropsychologia, 9(1), 97–113. doi:10.1016/0028-3932(71)90067-4

Pauletti, C., Mannarelli, D., Grippo, A., Currà, A., Locuratolo, N., De Lucia, M. C., & Fattapposta, F. (2014). Phasic alertness in a cued double-choice reaction time task: A contingent negative variation (CNV) study. Neuroscience Letters, 581, 7–13. doi:10.1016/j.neulet.2014.07.059

Polich, J. (2007). Updating P300: An integrative theory of P3a and P3b. Clinical Neurophysiology, 118(10), 2128–2148. doi:10.1016/j.clinph.2007.04.019

Polich, J., & Criado, J. R. (2006). Neuropsychology and neuropharmacology of P3a and P3b. International Journal of Psychophysiology, 60(2), 172–185. doi:10.1016/j.ijpsycho.2005.12.012

Romano, J. C., Howard, J. H., Jr., & Howard, D. V. (2010). One-year retention of general and sequence-specific skills in a probabilistic, serial reaction time task. Memory, 18(4), 427–441. doi:10.1080/09658211003742680

Rose, M., Verleger, R., & Wascher, E. (2001). ERP correlates of associative learning. Psychophysiology, 38(3), 440–450. doi:10.1111/1469-8986.3830440

Rüsseler, J., Hennighausen, E., Münte, T. F., & Rösler, F. (2003). Differences in incidental and intentional learning of sensorimotor sequences as revealed by event-related brain potentials. Cognitive Brain Research, 15(2), 116–126. doi:10.1016/S0926-6410(02)00145-3

Rüsseler, J., Münte, T. F., & Wiswede, D. (2018). On the influence of informational content and key-response effect mapping on implicit learning and error monitoring in the serial reaction time (SRT) task. Experimental Brain Research, 236(1), 259–273. doi:10.1007/s00221-017-5124-z

Rüsseler, J., & Rösler, F. (2000). Implicit and explicit learning of event sequences: Evidence for distinct coding of perceptual and motor representations. Acta Psychologica, 104(1), 45–67. doi:10.1016/S0001-6918(99)00053-0

Schumacher, E. H., Hendricks, M. J., & D’Esposito, M. (2005). Sustained involvement of a frontal-parietal network for spatial response selection with practice of a spatial choice-reaction task. Neuropsychologia, 43(10), 1444–1455. doi:10.1016/j.neuropsychologia.2005.01.002

Shohamy, D., & Daw, N. D. (2015). Integrating memories to guide decisions. Current Opinion in Behavioral Sciences, 5, 85–90. doi:10.1016/j.cobeha.2015.08.010

Simor, P., Zavecz, Z., Horváth, K., Éltető, N., Török, C., Pesthy, O., … Nemeth, D. (2019). Deconstructing procedural memory: Different learning trajectories and consolidation of sequence and statistical learning. Frontiers in Psychology, 9(2708). doi:10.3389/fpsyg.2018.02708

Song, S., Howard, J. H., Jr., & Howard, D. V. (2007). Sleep does not benefit probabilistic motor sequence learning. The Journal of Neuroscience, 27(46), 12475–12483. doi:10.1523/jneurosci.2062-07.2007

Stadler, W., Klimesch, W., Pouthas, V., & Ragot, R. (2006). Differential effects of the stimulus sequence on CNV and P300. Brain Research, 1123(1), 157–167. doi:10.1016/j.brainres.2006.09.040

Stark-Inbar, A., Raza, M., Taylor, J. A., & Ivry, R. B. (2017). Individual differences in implicit motor learning: Task specificity in sensorimotor adaptation and sequence learning. Journal of Neurophysiology, 117(1), 412–428. doi:10.1152/jn.01141.2015

Stock, A. K., Steenbergen, L., Colzato, L., & Beste, C. (2016). The system neurophysiological basis of non-adaptive cognitive control: Inhibition of implicit learning mediated by right prefrontal regions. Human Brain Mapping, 37(12), 4511–4522. doi:10.1002/hbm.23325

Szegedi-Hallgató, E., Janacsek, K., & Nemeth, D. (2019). Different levels of statistical learning - hidden potentials of sequence learning tasks. PLoS ONE, 14(9), e0221966. doi:10.1371/journal.pone.0221966

Szegedi-Hallgató, E., Janacsek, K., Vékony, T., Tasi, L. A., Kerepes, L., Hompoth, E. A., … Németh, D. (2017). Explicit instructions and consolidation promote rewiring of automatic behaviors in the human mind. Scientific Reports, 7(1), 4365. doi:10.1038/s41598-017-04500-3

Takács, Á., Kóbor, A., Chezan, J., Éltető, N., Tárnok, Z., Nemeth, D., … Janacsek, K. (2018). Is procedural memory enhanced in Tourette syndrome? Evidence from a sequence learning task. Cortex, 100, 84–94. doi:10.1016/j.cortex.2017.08.037

Takács, Á., Shilon, Y., Janacsek, K., Kóbor, A., Tremblay, A., Nemeth, D., & Ullman, M. T. (2017). Procedural learning in Tourette syndrome, ADHD, and comorbid Tourette-ADHD: Evidence from a probabilistic sequence learning task. Brain and Cognition, 117, 33–40. doi:10.1016/j.bandc.2017.06.009

Tóth-Fáber, E., Janacsek, K., Szőllősi, Á., Kéri, S., & Németh, D. (2020). Regularity extraction under stress: Boosted statistical learning but unaffected sequence learning. bioRxiv, 2020.2005.2013.092726. doi:10.1101/2020.05.13.092726

Tóth, B., Janacsek, K., Takács, Á., Kóbor, A., Zavecz, Z., & Nemeth, D. (2017). Dynamics of EEG functional connectivity during statistical learning. Neurobiology of Learning and Memory, 144, 216–229. doi:10.1016/j.nlm.2017.07.015

Török, B., Janacsek, K., Nagy, D. G., Orbán, G., & Nemeth, D. (2017). Measuring and filtering reactive inhibition is essential for assessing serial decision making and learning. Journal of Experimental Psychology: General, 146(4), 529–542. doi:10.1037/xge0000288

Tremblay, S., & Saint-Aubin, J. (2009). Evidence of anticipatory eye movements in the spatial Hebb repetition effect: Insights for modeling sequence learning. Journal of Experimental Psychology: Learning, Memory, and Cognition, 35(5), 1256–1265. doi:10.1037/a0016566

Twomey, D. M., Murphy, P. R., Kelly, S. P., & O’Connell, R. G. (2015). The classic P300 encodes a build-to-threshold decision variable. European Journal of Neuroscience, 42(1), 1636–1643. doi:10.1111/ejn.12936

van Boxtel, G. J. M., & Böcker, K. B. E. (2004). Cortical measures of anticipation. Journal of Psychophysiology, 18(2-3), 61–76. doi:10.1027/0269-8803.18.23.61

van Boxtel, G. J. M., & Brunia, C. H. M. (1994). Motor and non-motor aspects of slow brain potentials. Biological Psychology, 38(1), 37–51. doi:10.1016/0301-0511(94)90048-5

Verleger, R., Hamann, L. M., Asanowicz, D., & Śmigasiewicz, K. (2015). Testing the S–R link hypothesis of P3b: The oddball effect on S1-evoked P3 gets reduced by increased task relevance of S2. Biological Psychology, 108, 25–35. doi:10.1016/j.biopsycho.2015.02.010

Verleger, R., Jaśkowski, P., & Wascher, E. (2005). Evidence for an integrative role of P3b in linking reaction to perception. Journal of Psychophysiology, 19(3), 165–181. doi:10.1027/0269-8803.19.3.165

Verleger, R., Paehge, T., Kolev, V., Yordanova, J., & Jaśkowski, P. (2006). On the relation of movement-related potentials to the go/no-go effect on P3. Biological Psychology, 73(3), 298–313. doi:10.1016/j.biopsycho.2006.05.005

Verleger, R., Paulick, C., Möcks, J., Smith, J. L., & Keller, K. (2013). Parafac and go/no-go: Disentangling CNV return from the P3 complex by trilinear component analysis. International Journal of Psychophysiology, 87(3), 289–300. doi:10.1016/j.ijpsycho.2012.08.003

Verleger, R., Siller, B., Ouyang, G., & Śmigasiewicz, K. (2017). Effects on P3 of spreading targets and response prompts apart. Biological Psychology, 126, 1–11. doi:10.1016/j.biopsycho.2017.03.011

Verleger, R., & Śmigasiewicz, K. (2016). Do rare stimuli evoke large P3s by being unexpected? A comparison of oddball effects between standard-oddball and prediction-oddball tasks. Advances in Cognitive Psychology, 12(2), 88–104. doi:10.5709/acp-0189-9

Verleger, R., Wauschkuhn, B., van der Lubbe, R., Jaśkowski, P., & Trillenberg, P. (2000). Posterior and anterior contribution of hand-movement preparation to late CNV. Journal of Psychophysiology, 14(2), 69–86. doi:10.1027//0269-8803.14.2.69

Virag, M., Janacsek, K., Horvath, A., Bujdoso, Z., Fabo, D., & Nemeth, D. (2015). Competition between frontal lobe functions and implicit sequence learning: Evidence from the long-term effects of alcohol. Experimental Brain Research, 233(7), 2081–2089. doi:10.1007/s00221-015-4279-8

Walter, W. G., Cooper, R., Aldridge, V. J., McCallum, W. C., & Winter, A. L. (1964). Contingent negative variation : An electric sign of sensori-motor association and expectancy in the human brain. Nature, 203(4943), 380–384. doi:10.1038/203380a0

Weerts, T. C., & Lang, P. J. (1973). The effects of eye fixation and stimulus and response location on the contingent negative variation (CNV). Biological Psychology, 1(1), 1–19. doi:10.1016/0301-0511(73)90010-0

Willingham, D. B., Greenberg, A. R., & Thomas, R. C. (1997). Response-to-stimulus interval does not affect implicit motor sequence learning, but does affect performance. Memory & Cognition, 25(4), 534–542. doi:10.3758/BF03201128

