## Supplementary Materials for "Implicit anticipation of probabilistic regularities: Larger CNV emerges for unpredictable events"

**Analysis of the response-locked P3 component**

**1. EEG/ERP analysis**

Up to the second step of segmentation, all preprocessing procedures were the same as described in the main text. Then, within each period, segments were extracted from -1400 ms to 400 ms relative to response onset, separately for pattern high-probability and random low-probability triplets (without trills and repetitions). Only those correctly responded triplets were included that also followed correctly responded triplets. Next, the response-locked segments were average baseline corrected from -1400 ms to 400 ms relative to response onset (Killikelly & Szűcs, 2013).^[1]^ After that, the same automatic artifact rejection algorithm was applied as in the case of the cue-locked segments with the further manual rejection of those segments that contained low frequency electrode drifts. The mean numbers of the retained (artifact-free) segments were 339.9 (*SD* = 39.9, range: 195 – 384) for pattern high-probability triplets and 168.8 (*SD* = 24.0, range: 78 – 211) for random low-probability triplets. Finally, these retained segments were averaged for the two triplet types in each of the three periods. The P3 peak in response-locked averages was quantified as the mean amplitude within 50 ms before to 50 ms after response onset where the grand average peak appeared over the centroparietal electrode pool. The late P3 in response-locked averages was quantified as the mean amplitude between 50 ms and 250 ms after response onset, since a negative deflection started on the grand average waveform at approximately 250 ms over the centroparietal electrode pool (cf. Kóbor et al., 2019). The statistical analysis of these mean amplitudes was performed as described in the main text.

**2. Results**

Grand average response-locked ERP waveforms for the two triplet types averaged for all periods and split by period over the centroparietal electrode pool are presented in Figures S1-S2, respectively. Note that the CNV is also apparent in the response-locked averages: It rises from around -600 ms (relative to response onset), develops until approximate target onset, and returns to baseline before response onset.

**2.1 P3 peak**

From the Type by Period ANOVA on the response-locked P3 peak, the main effect of Type was a tendency, *F*(1, 35) = 2.97, *p* = .094, η*_p_*^2^ = .078, indicating that the P3 peak amplitude tended to be lower for random low-probability triplets than for pattern high-probability ones (3.41 µV vs. 3.46 µV; see Fig. S1B, C, Fig. S3A). The Period main effect, *F*(2, 70) = 0.77, ε = .766, *p* = .438, η*_p_*^2^ = .021, and the Type * Period interaction, *F*(2, 70) = 1.82, *p* = .169, η*_p_*^2^ = .050, were not significant (see Fig. S2 and Fig. S3A). When the same Type by Period ANOVA was performed on the response-locked P3 peak measured only at electrode Pz or over a parietal electrode pool (average activity of electrodes CPz, CP1, CP2, Pz, P1, and P2), as in our previous study (Kóbor et al., 2019), the main effect of Type was significant (Pz: *F*(1, 35) = 6.47, *p* = .016, η*_p_*^2^ = .156, 4.13 µV vs. 4.22 µV; parietal pool: *F*(1, 35) = 6.33, *p* = .017, η*_p_*^2^ = .153, 3.95 µV vs. 4.03 µV, see Fig. S1A).

**2.2 Late P3**

Type by Period ANOVA on the response-locked late P3 revealed a significant main effect of Type, *F*(1, 35) = 10.41, *p* = .003, η*_p_*^2^ = .229, indicating that the mean amplitude of the late P3 was lower for random low-probability triplets than for pattern high-probability ones (1.51 µV vs. 1.59 µV; see Fig. S1B, C, Fig. S3B). The Period main effect, *F*(2, 70) = 0.11, ε = .761, *p* = .840, η*_p_*^2^ = .003, and the Type * Period interaction, *F*(2, 70) = 0.20, *p* = .821, η*_p_*^2^ = .006, were not significant (see Fig. S2 and Fig. S3B). As in the case of the P3 peak, the Type main effect was significant when the response-locked late P3 was measured at electrode Pz or over a parietal electrode pool (Pz: *F*(1, 35) = 5.57, *p* = .024, η*_p_*^2^ = .137, 0.47 µV vs. 0.57 µV; parietal pool: *F*(1, 35) = 8.78, *p* = .005, η*_p_*^2^ = .200, 1.08 µV vs. 1.18 µV, see Fig. S1A).

**2.3 Correlations between the behavioral and the P3 measures**

The response-locked mean amplitudes measured over the centroparietal electrode pool were included in the correlational analyses. Significant negative correlations emerged between RTs and the P3 peak in the first (*r*(34) = -.334, *p* = .046) and second (*r*(34) = -.360, *p* = .031) periods for pattern high-probability triplets and in the first period (*r*(34) = -.342, *p* = .041) for random low-probability triplets. Significant negative correlations emerged also between accuracy and the P3 peak in the first (*r*(34) = -.372, *p* = .025) and second (*r*(34) = -.351, *p* = .036) periods for pattern high-probability triplets and in the last period (*r*(34) = -.334, *p* = .047) for random low-probability triplets. Thus, some indication was found that faster and less accurate responses were associated with larger P3 peak amplitudes. No significant correlations were found between the behavioral measures and the late P3 (all |*r*|s ≤ .212, *p*s ≥ .215).

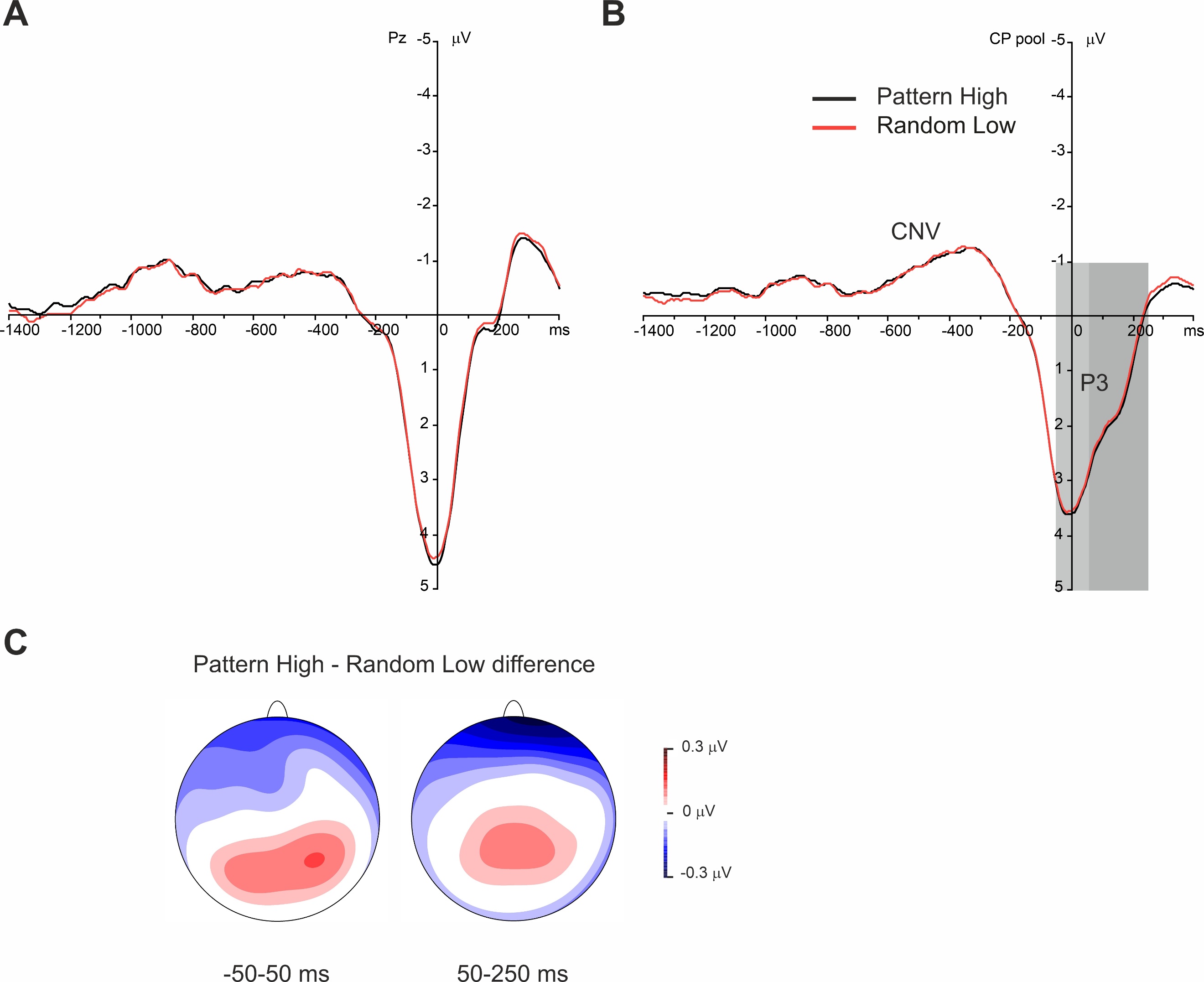

**Figure S1.** Grand average response-locked ERP waveforms at (A) electrode Pz and over (B) the centroparietal electrode pool are presented, displaying the P3 component for pattern high-probability and random low-probability triplets, averaged for all periods. Zero ms denotes response onset. In part (B) of this figure, the medium-grey shaded area indicates the time window in which the P3 peak was quantified (-50-50 ms), and the dark-grey shaded area indicates the time window in which the late P3 was quantified (50-250 ms). Negativity is plotted upwards here and in Figure S2. (C) The scalp topography (amplitude distribution) of ERP differences for pattern high-probability minus random low-probability triplets in the time windows of the P3 peak (left) and the late P3 (right), averaged for all periods.

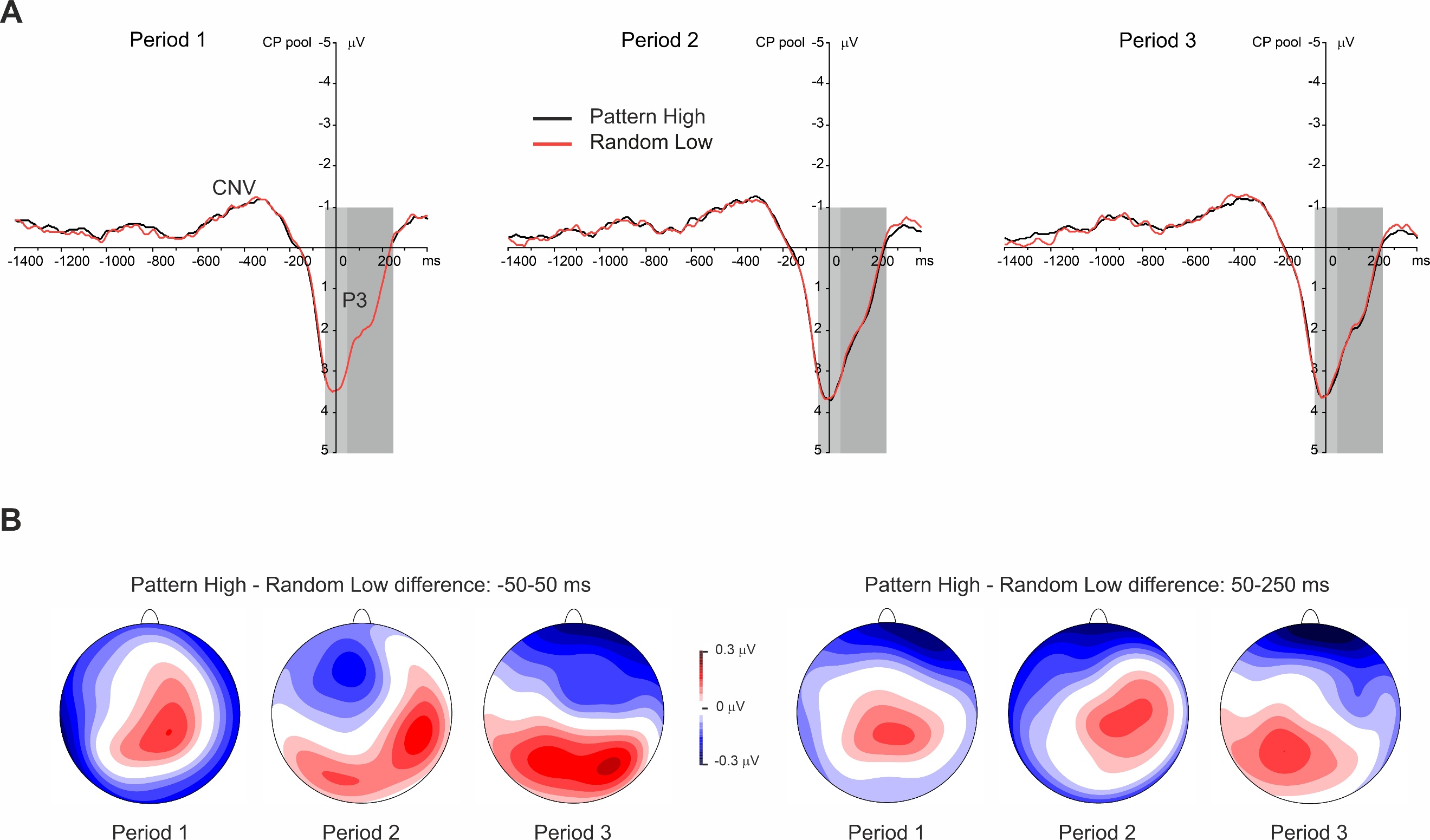

**Figure S2.** (A) Grand average response-locked ERP waveforms over the centroparietal electrode pool are presented, displaying the P3 component for each period (1–3) and triplet type (pattern high-probability and random low-probability triplets). Zero ms denotes response onset. The medium-grey shaded area is the time window of the P3 peak (-50-50 ms), and the dark-grey shaded area is the time window of the late P3 (50-250 ms). (B) The scalp topography (amplitude distribution) of ERP differences in each period for pattern high-probability minus random low-probability triplets in the time windows of the P3 peak (left) and the late P3 (right).

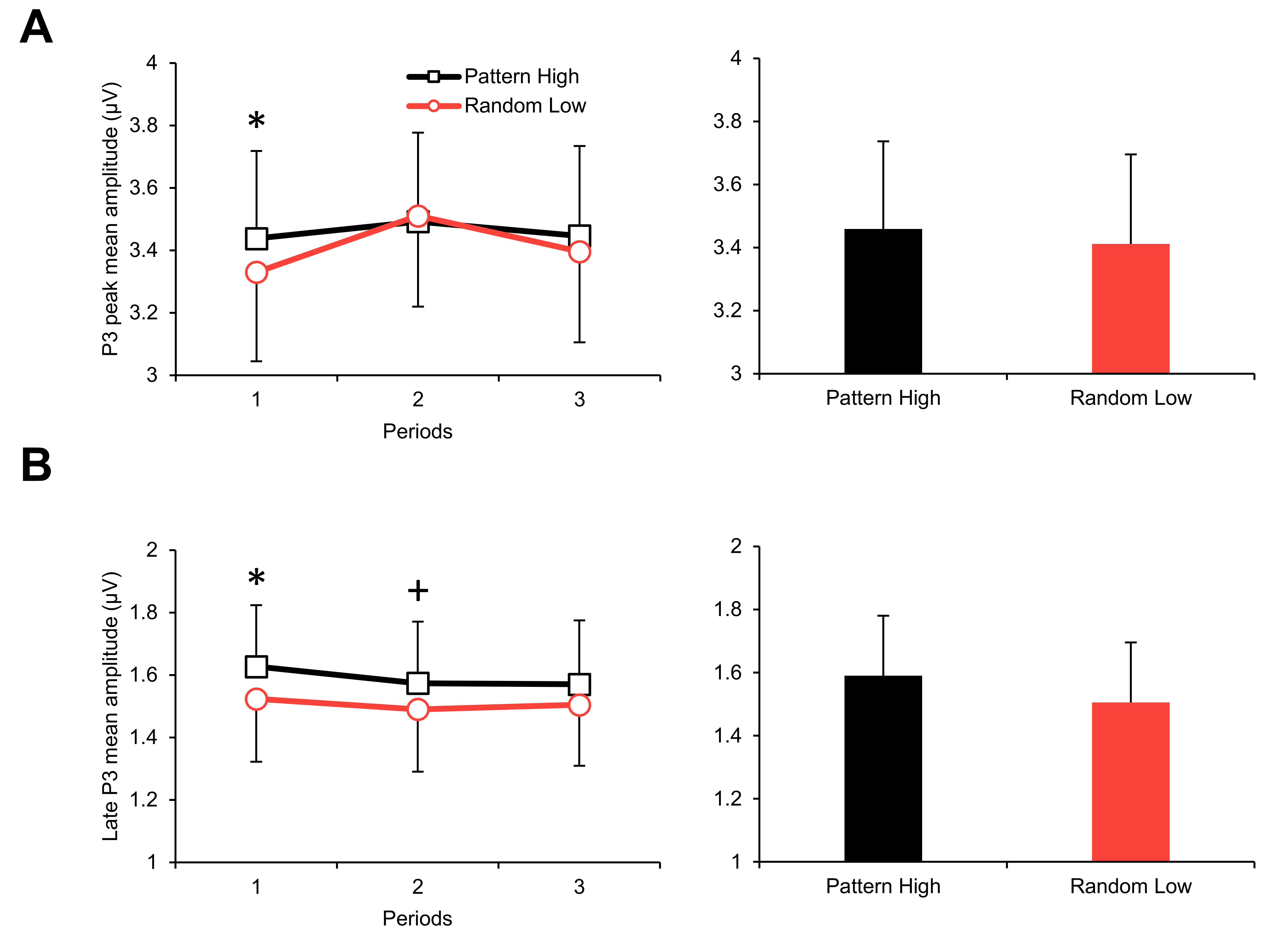

**Figure S3.** (A) Group-average response-locked P3 peak mean amplitudes split by period (1–3) and triplet type (pattern high-probability vs. random low-probability triplets) (left panel, line chart) and split by triplet type but collapsed across periods (right panel, bar chart). (B) Group-average response-locked late P3 mean amplitudes split by period and triplet type (left panel, line chart) and split by triplet type but collapsed across periods (right panel, bar chart). Error bars denote standard error of mean. On the left panel, asterisks/plus marks above the means of each period denote the significance of the difference between triplet types (note that these pairwise differences are indicated although nonsignificant Type * Period interactions were found).

*Note*: + *p* < .010, * *p* < .050

**References**

Killikelly, C., & Szűcs, D. (2013). Delayed development of proactive response preparation in adolescents: ERP and EMG evidence. *Developmental Cognitive Neuroscience, 3*, 33-43. doi:10.1016/j.dcn.2012.08.002

Kóbor, A., Horváth, K., Kardos, Z., Takács, Á., Janacsek, K., Csépe, V., & Nemeth, D. (2019). Tracking the implicit acquisition of nonadjacent transitional probabilities by ERPs. *Memory & Cognition, 47*(8), 1546-1566. doi:10.3758/s13421-019-00949-x

1. The -1400 ms to 400 ms segment length was chosen to include most of the response preparation and execution phases, from cue onset until target presentation. Particularly, the mean RT was 369 ms (*SD* = 66 ms) and 95% of all responses were equal or below 485 ms for the two triplet types. Mean activity of the entire segment as the baseline was chosen since no obvious neutral time interval occurred in the segment and the inter-trial-interval was 1800 ms.
